# Volume-transmitted GABA waves pace epileptiform rhythms in the hippocampal network

**DOI:** 10.1101/2021.03.25.437016

**Authors:** Vincent Magloire, Leonid P. Savtchenko, Sergyi Sylantyev, Thomas P. Jensen, Nicholas Cole, Jonathan S. Marvin, Loren L. Looger, Dimitri M. Kullmann, Matthew C. Walker, Ivan Pavlov, Dmitri A. Rusakov

## Abstract

Mechanisms that entrain and drive rhythmic epileptiform discharges remain debated. Traditionally, this quest has been focusing on interneuronal networks driven by GABAergic connections that activate synaptic or extrasynaptic receptors. However, synchronised interneuronal discharges could also trigger a transient elevation of extracellular GABA across the tissue volume, thus raising tonic GABA_A_ receptor conductance (*G*_tonic_) in multiple cells. Here, we use patch-clamp GABA ‘sniffer’ and optical GABA sensor to show that periodic epileptiform discharges are preceded by region-wide, rising waves of extracellular GABA. Neural network simulations that incorporate volume-transmitted GABA signals point to mechanistic principles underpinning this relationship. We validate this hypothesis using simultaneous patch-clamp recordings from multiple nerve cells, selective optogenetic stimulation of fast-spiking interneurons. Critically, we manipulate GABA uptake to suppress extracellular GABA waves but not synaptic GABAergic transmission, which shows a clear effect on rhythm generation. Our findings thus unveil a key role of extrasynaptic, volume-transmitted GABA actions in pacing regenerative rhythmic activity in brain networks.

## INTRODUCTION

GABA is the principal inhibitory neurotransmitter in the forebrain. It mediates conventional fast synaptic transmission through GABA_A_ receptors (GABA_A_Rs), as well as slower synaptic transmission mediated by GABA_B_ receptors. There is also a slow GABA_A_R-mediated conductance (*G*_tonic_), often referred to as ‘tonic inhibition’, which reflects activation of slowly desensitizing, extrasynaptic GABA_A_Rs by GABA that is present throughout the extracellular space. Historically, micro-dialysis studies *in vivo* have suggested that the ambient extracellular GABA concentration ([GABA]_e_) is relatively stable, even though it may depend on the physiological or behavioural state (van der Zeyden et al., 2008). Indeed, long-lasting changes in [GABA]_e_ due to extrasynaptic GABA escape should be constrained by GABA transporters (Scanziani, 2000; Overstreet and Westbrook, 2003). However, the dialysis technique usually reports [GABA]_e_ averaged over tens of seconds whereas the relatively low apparent affinity (*K_m_* = 10-20 μM) of the main GABA transporter GAT-1 (Loo et al., 2000; MacAulay et al., 2003; Bicho and Grewer, 2005) cannot prevent faster, low micromolar-range activity-dependent fluctuations in [GABA]_e_.

The dynamics of *G*_tonic_ has been associated with the firing pattern of interneurons (Glykys and Mody, 2007) and implicated in epileptiform discharge initiation (Scanziani, 2000; Marchionni and Maccaferri, 2009). Activation of extrasynaptic GABA_A_Rs can also alter the excitability of principal cells (Brickley et al., 2001; Scimemi et al., 2005a; Pavlov et al., 2009; Song et al., 2011), which directly influences network behaviour (Mann and Mody, 2010; Pavlov et al., 2014) including input coincidence detection rules (Pouille and Scanziani, 2001; Pavlov et al., 2011; Sylantyev et al., 2020). Whilst a collapse in inhibitory activity has been associated with the emergence of epileptiform events (Karlocai et al., 2014), hippocampal interneurons appear to maintain or increase their firing rate before the onset of ictal activity (Miri et al., 2018). In the cortex, seizure-like events in pyramidal neurons can be preceded by an outward GABA_A_R current and rises in extracellular K^+^ (Librizzi et al., 2017). Furthermore, optogenetic activation of GABAergic cells *en masse* can paradoxically trigger ictal events, possibly as a result of post-inhibitory rebound excitation (Chang et al., 2018). While these observations clearly implicate dynamic interactions between network inhibition and disinhibition (or rebound excitation) in the generation of ictal discharges, the precise nature of these interactions in regulating rhythmic epileptiform activity remain poorly understood.

Intriguingly, we have previously found a biphasic, bell-shaped relationship between *G*_tonic_ (controlled through either dynamic clamp or by changing ambient [GABA]_e_), and the firing and synchronisation of interneuronal networks (Song et al., 2011; Pavlov et al., 2014). Because *G*_tonic_ itself depends on interneuronal firing, this type of relationship suggests, at least in theory, an inherent capacity for self-organising rhythms, or periodic waves of activity (Huguenard and McCormick, 1992). Genetically encoded optical GABA sensors have recently enabled the detection of prominent [GABA]_e_ transients (on a 10-100 ms timescale) during synchronous ictal discharges *in vivo* (Marvin et al., 2019), arguing that fluctuations in *G*_tonic_ may have a role in periodic network activity in the intact brain.

We therefore sought to understand how the dynamics of [GABA]_e_ and *G*_tonic_ relate to the rhythmic activity of neuronal networks. To this end, we focused on FS hippocampal interneurons hypothesising that their synchronized firing should drive transient volume-transmitted changes in [GABA]_e_ that regulate network rhythms, such as interictal epileptiform discharges. We tested this hypothesis by combining outside-out membrane patch (GABA sniffer) recordings (Isaacson et al., 1993; Wlodarczyk et al., 2013), an optical GABA sensor iGABASnFR (Marvin et al., 2019), and optogenetic stimulation of PV-positive interneurons, with computer simulations of spiking neural networks (Savtchenko and Rusakov, 2014; Aleksin et al., 2017). Importantly, manipulating GABA uptake to moderate extracellular GABA waves but not synaptic GABAergic transmission, shows a clear effect on rhythm generation.

## RESULTS

### Transient [GABA]_e_ rises in tissue tend to precede interictal events

First, to examine the relationship between regular bursts of neuronal firing and [GABA]_e_, we used acute hippocampal slices perfused with a solution containing 0 [Mg^2+^] and 5 mM [K^+^], in order to elicit epileptiform activity. In the CA1 region, this activity was detected as brief (<200 ms) periodic field-potential spikes that reflected interictal bursts of neuronal populations (Figure 1A, top trace). In parallel, we monitored local [GABA]_e_ using an outside-out ‘sniffer’ patch which reports GABA_A_R channel activity (Isaacson et al., 1993), as previously described (Wlodarczyk et al., 2013; Sylantyev et al., 2020) (Figure 1A, middle; Methods). Channel activity bursts in the patch appeared synchronised with interictal discharges (Figure 1A), but a detailed analysis revealed that channel activity always started to rise 0.5-1 s before interictal spikes (Figure 1B-C; Figure S1A; summary in Figure 1D). We took special care to prepare uniformly shaped sniffer patch pipettes for these experiments, and used a multi-channel rapid solution exchange to calibrate [GABA]_e_ against recorded GABA_A_R channel activity (Sylantyev and Rusakov, 2013a), giving EC_50_ value of ~1.1 μM, which corresponds to a charge transfer rate (or current) of 256 ± 57 fC·s^-1^ (fA; n = 6; Figure 1E). Based on this, the present recordings indicated that [GABA]_e_ rises from its resting level of ~0.3 μM to a peak of 1.5-2 μM during interictal discharges (Figure 1E).

**Figure 1.**
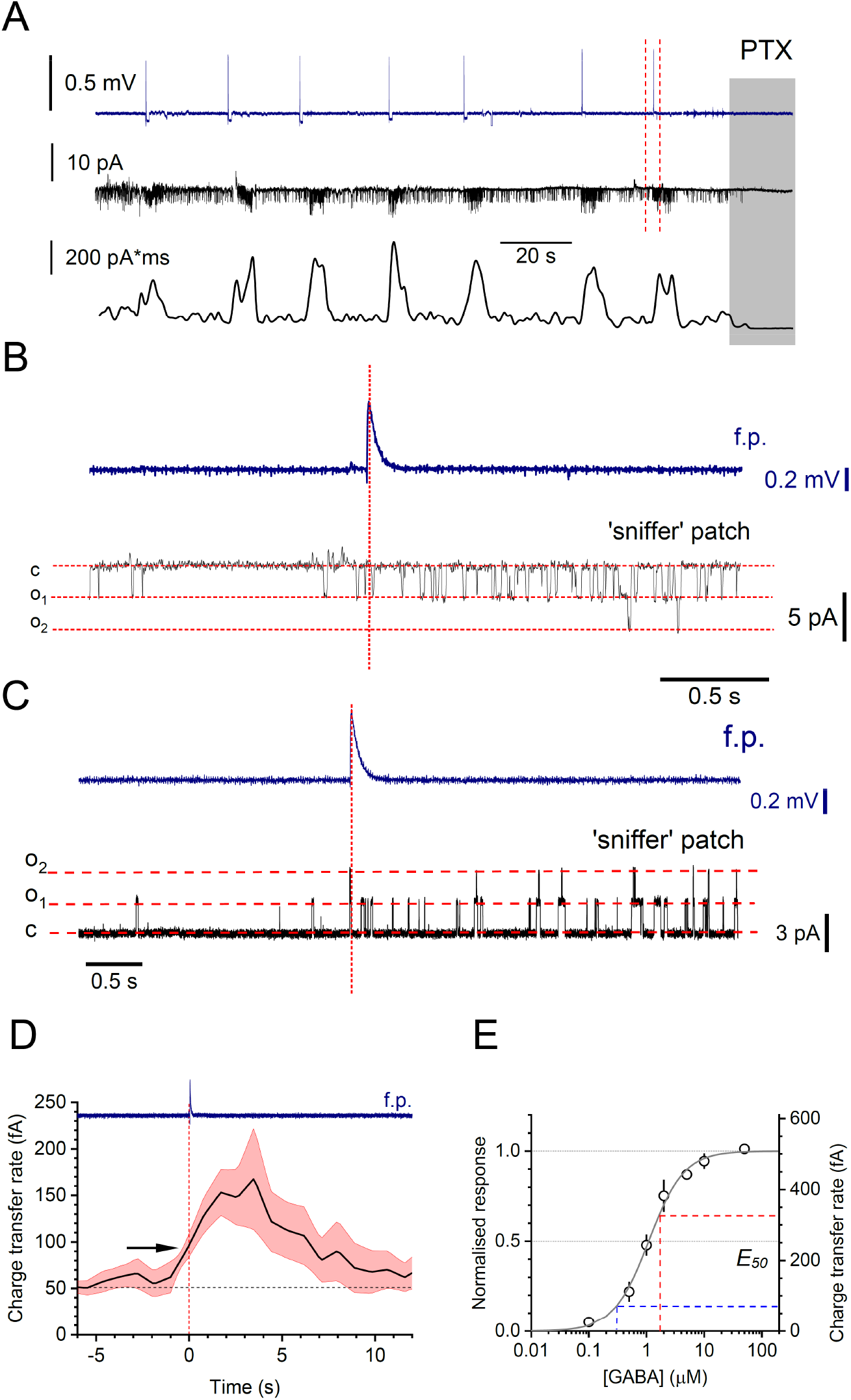
Rhythmic fluctuations of extracellular GABA during interictal activity in the hippocampal area CA1. (**A**) Local field potentials (LFP, top trace), sniffer-patch single-channel activity (middle), and charge dynamics (bottom) during interictal discharges in hippocampal slices; grey shade, picrotoxin (PTX) application at the end of trials. (**B**) Fragment in A (between dotted lines) expanded to show the LFP (top) and patch recording (black) details; high chloride pipette solution was used to record inward GABA_A_R-mediated channel openings (V_m_ = −70 mV). (**C**) Similar setting to B, but with low chloride solution to document outward currents (V_m_ = 0 mV). (**D**) Averaged time course of charge transfer rate (equal to current; mean ± s.e.m. for slice-average traces, n = 6 slices) at the onset of individual f.p. (vertical dashed line, blue trace); arrow, interictal discharge onset coincides with GABA_A_R activity reaching ~35% of its peak. (**E**) Normalised and absolute charge transfer values recorded in sniffer patches at different [GABA]_e_ (dots, mean ± s.e.m.; n = 4-6 for individual values; grey line, sigmoid best-fit); dashed lines, [GABA]_e_ during resting level (blue) and peak (red) of channel activity recorded.

We next explored the cellular basis of these observations. The spiking activity of individual FS PV+ interneurons increased substantially from their basal level several seconds before the peak of CA1 pyramidal cell discharges (which correspond to the interictal events, Figure S1A-B). This was faithfully reflected by the elevated inhibitory input to CA1 pyramidal cells before the interictal bursts (Figure S1B-C). In contrast, pyramidal cell spiking appeared time-locked to the epileptiform discharges (Figure S1D-E). These observations reflect earlier findings that interictal bursting is preceded by increased firing of FS PV+ interneurons (Karlocai et al., 2014; de Curtis and Avoli, 2016; Librizzi et al., 2017; Chang et al., 2018), which we suggest here to represent a ‘global’ wave of [GABA]_e_ detected with the ‘sniffer’ patch (Figure 1), rather than synaptic feedforward inhibition per se.

### Interictal spikes are preceded by [GABA]_e_ elevations in the pyramidal cell layer

Whilst the ‘sniffer’ patch can detect [GABA]_e_ changes with high sensitivity, it reports average [GABA]_e_ values near the slice surface, potentially far away from the sites of GABA release. To understand whether [GABA]_e_ displays similar dynamics within the pyramidal cell layer (one of the main axon target areas of FS interneurons), deep within the tissue, we used the recently developed optical GABA sensor iGABASnFR (Marvin et al., 2019). We expressed the sensor in a proportion of CA1 and CA3 pyramidal cells under the *hSynapsin-1* promoter, in organotypic hippocampal slices (Figure 2A; Methods): sparse expression of iGABASnFR helped to minimise any potential buffering effects on the dynamics of extracellular GABA (Armbruster et al., 2020; Kopach et al., 2020).

**Figure 2.**
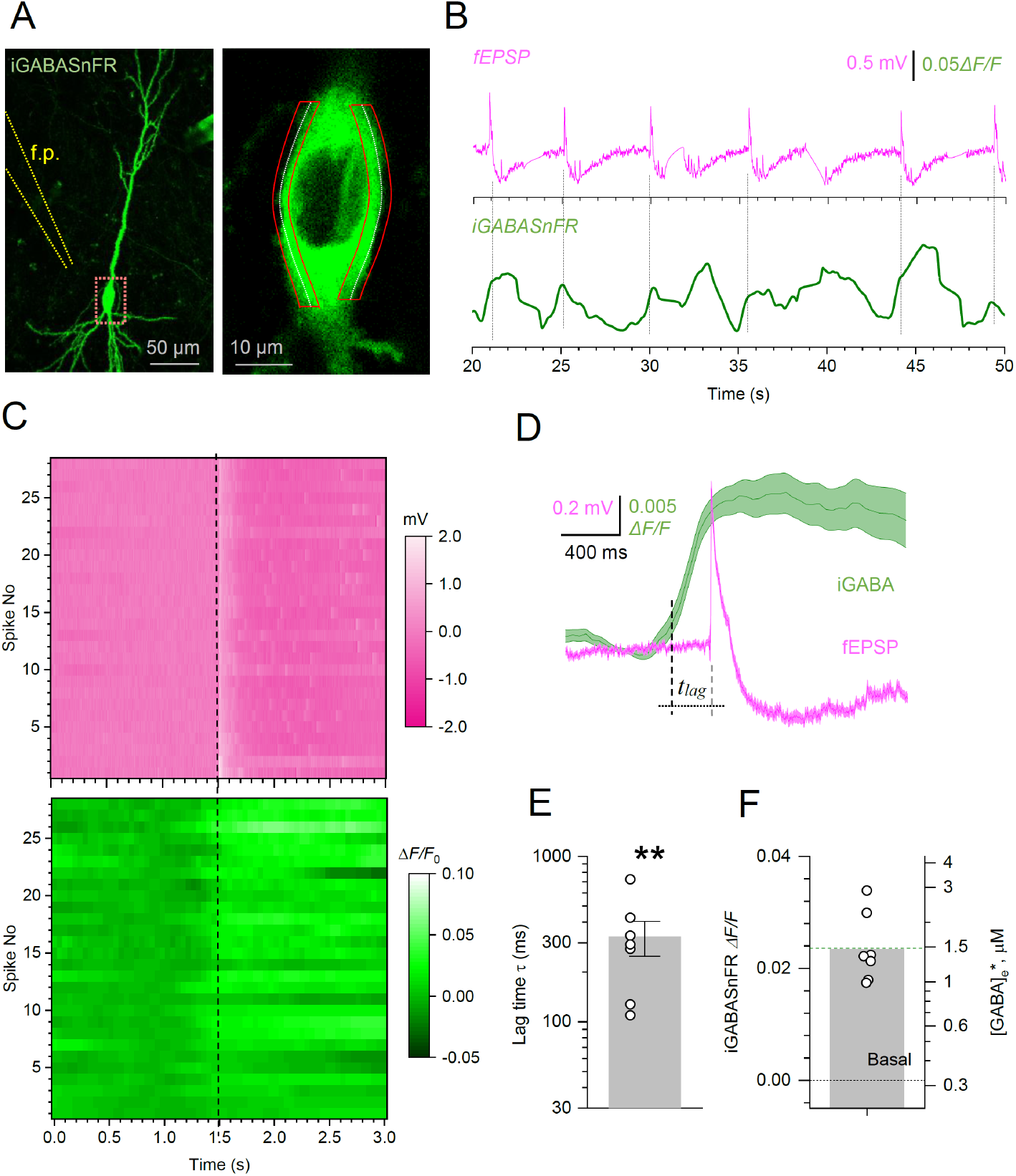
Optical registration of [GABA]_e_ waves during interictal events in hippocampal slices. (**A**) CA1 pyramidal cell expressing iGABASnFR with a field pipette electrode (f.p., left) and two imaging ROIs near the cell soma periphery (right, zoomed-in fragment shown by red rectangle on the left; λ_x_^2p^ = 910 nm). (**B**) *O*ne-slice example, fEPSPs (top) and iGABASnFR signal (bottom) recorded during interictal spikes. (**C**) Diagram showing 28 interictal fEPSPs aligned with respect to their peak times (top), and the corresponding iGABASnFR signals (bottom; false colour scale; one-slice example). (**D**) Summary of recordings shown in C, at expanded time scale: traces, mean ± s.e.m. (n = 28); dotted lines, lag time <u>*t_lag_*</u> between the 20% peak rise time of iGABASnFR signal and peak time of fEPSP, as indicated. (**E**) Summary for *τ* values obtained in seven slices (mean ± s.e.m.; dots, data from individual slices; ** *p* = 0.0052, paired Student’s *t*-test). (**F**) Average peak iGABASnFR signal *ΔF/F* values for seven experiments as in E (left ordinate; column, mean), and the corresponding estimated [GABA] values ([GABA]_e_*, right ordinate).

We next imaged iGABASnFR fluorescence within microscopic regions of interest (ROIs) near the periphery of individual pyramidal cell somata using two-photon excitation microscopy, as described earlier (Jensen et al., 2019; Henneberger et al., 2020) while recording field potentials nearby (Figure 2A). Periodic interictal events were consistently accompanied by transient elevations of iGABASnFR fluorescence that, again, appeared to be rising before individual field discharges (Figure 2B). The analysis of field potentials and iGABASnFR signals at higher resolution confirmed this trend (Figure 2C): the average interval from the iGABASnFR signal rise onset (taken at 20% signal peak, for conservative estimate) to the field potential spike onset was 327 ± 77 ms (mean ± s.e.m., n = 7 slices; Figure 2C-E; Figure S2). This time lag was orders of magnitude greater than the lag from a hypothetical pacemaker elsewhere in the hippocampal slice, arguing against the hypothesis that iGABASnFR is merely detecting a barrage of feed-forward inhibition.

During interictal cycles, the iGABASnFR signal peaked at 0.024 ± 0.002 *ΔF/F* (mean ± s.e.m.; Figure 2F, left ordinate). Membrane-bound iGABASnFR expressed extracellularly in hippocampal neurons and calibrated using clamped [GABA]_e_, shows an affinity of ~30 μM, with a saturating signal of ~0.7 *ΔF/F* (Marvin et al., 2019). In accordance with the Hill equation, and with resting [GABA]_e_ ~0.3 μM (Figure 1E), this suggests that 0.024 *ΔF/F* signal corresponds to a ~1.2 μM rise in [GABA]_e_ (Figure 2F, right ordinate), in excellent agreement with the ‘sniffer’ patch data (Figure 1E). We note that the [GABA]_e_ transients reported with iGABASnFR are subject to greater fluctuations than sniffer patch recordings, most likely because the latter represent [GABA]_e_ integrated over much larger tissue volumes.

### Biophysical basis of [GABA]_e_ -dependent network rhythms

To understand the role of [GABA]_e_ fluctuations in the generation of periodic interictal discharges, we simulated a network of FS hippocampal interneurons, with a stochastic excitatory input to individual cells (Figure 3A; Methods), by building upon our previous network models (Pavlov et al., 2014; Savtchenko and Rusakov, 2014; Aleksin et al., 2017). Several elegant modelling studies have provided plausible mechanistic explanations for the network regimes generating high-frequency rhythms including sharp-wave ripples (Ferguson et al., 2013; Schlingloff et al., 2014; Rich et al., 2019). However, the mechanisms proposed previously considered neither the 0.2-1.0 Hz frequency range, which is relevant here, nor the potential role of volume-transmitted GABA. An early theoretical study suggested a volume-transmitted feedback mechanism involving K^+^ waves as the main drive of 2-5 Hz network oscillations (Ho and Truccolo, 2016), but the effect of K^+^ fluctuations on glutamate and GABA uptake remained to be ascertained. Here, the key hypothesis was that interneuronal discharges generated both the IPSCs in individual cells and volume-transmitted [GABA]_e_ rises (Figure 3A) that contributed to *G*_tonic_ across the cell population (Savtchenko and Rusakov, 2014; Aleksin et al., 2017). *G*_tonic_ was thus set as a monotonic function of [GABA]_e_ (Methods), which in turn was determined by (i) a scalable [GABA]_e_ increment upon each individual synaptic discharge, and (ii) extracellular GABA removal by the main GABA transporter GAT-1, yielding an exponential [GABA]_e_ decay (Bicho and Grewer, 2005; Savtchenko et al., 2015a) (Methods).

**Figure 3.**
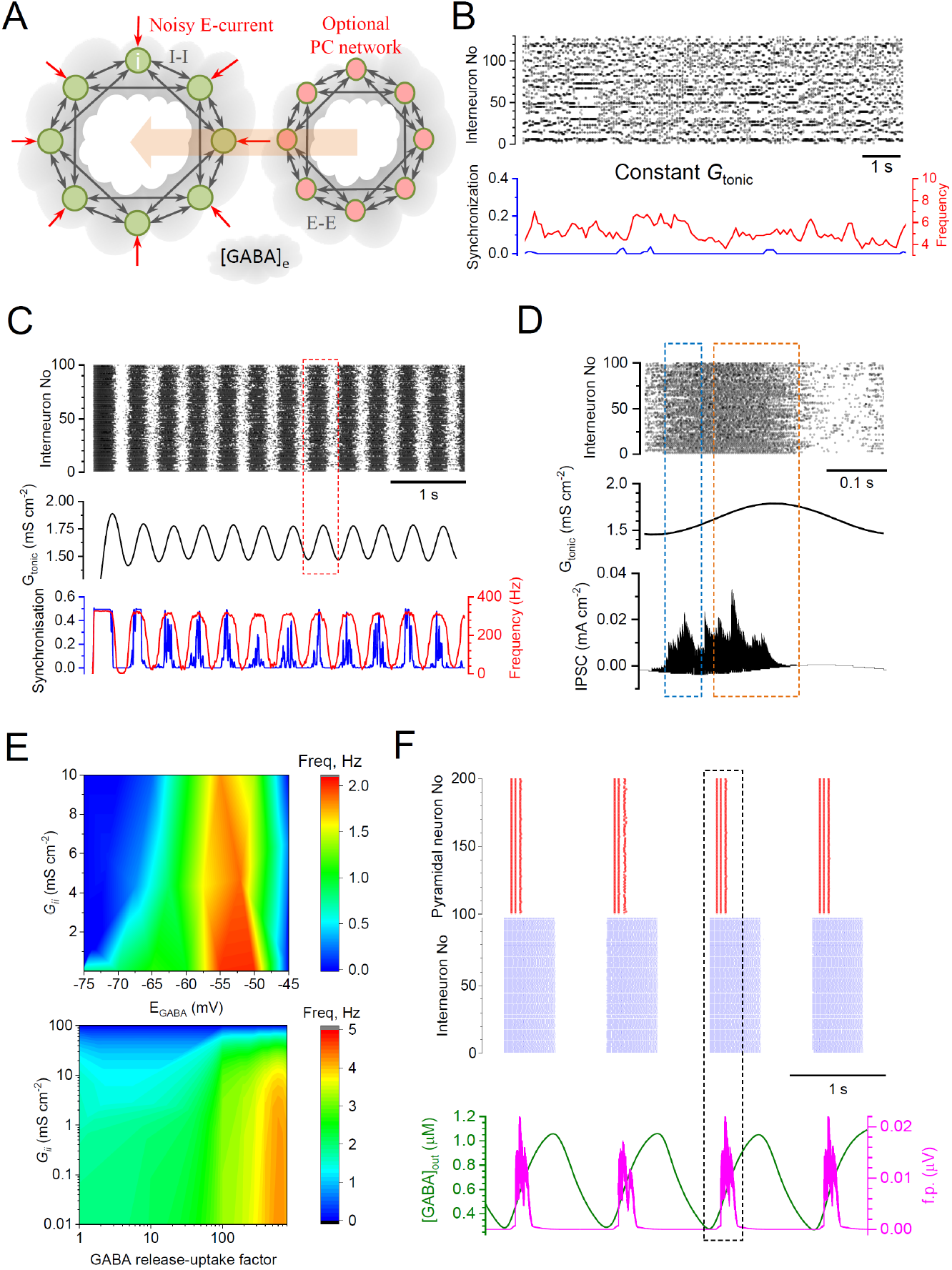
[GABA]_e_-dependent *G*_tonic_ robustly drives rhythmic activity of a modelled interneuron network across a range of parameters. (**A**) Schematic, FS interneuron network model (*I-I*, inhibitory synaptic connections), with external excitatory inputs (*E*-current), volume-transmitted [GABA]_e_ signal generating *G*_tonic_, and an optional network of pyramidal cells incorporated (*E-E*, excitatory connections). (**B**) Raster plot of interneuronal network spiking (top), network synchronisation coefficient (bottom, blue), and time course of average spiking frequency (bottom, red) for simulated network with constant *G*_tonic_ value (0.44 mS cm^-2^). (**C**) Raster plot as in B but with *G*_tonic_ driven by [GABA]_e_ (middle) calculated from integrated interneuronal discharges; network synchrony and mean network frequency show clear periodicity (bottom). Key model parameters: cell number *N* = 100, intra-network peak synaptic conductance *G*_ii_ = 0.1 nS cm^-2^; *E*-currents (Poisson series) with average synaptic conductance *g*_s_=0.02 mS cm^-2^, decay constant tau = 3 ms, and frequency *f_s_* = 100 Hz; GABA_A_R reversal potential *V_GABA_* = −56 mV, GABA release factor *A_f_* = 1.35 ×10^-7^ nS cm^-2^ ms^-1^, *G*_pump_ =0.003 s^-1^ (Methods). (**D**) Fragment from C (red dotted rectangle) enlarged, with the integrated IPSC time course (bottom). Blue and orange dotted rectangles indicate high-frequency non-synchronised, and oscillating and synchronised IPSC periods, respectively. (**E**) Parameter-space heatmaps: rhythm generation over a range of *G_ii_* and *E_GABA_* (*R* = 1, top), and *G_ii_* and release-uptake factor *R* (Methods; *E_GABA_* = −65 mV, bottom); deep blue area indicates no detectable rhythmic activity. (**F**) *Top:* Raster plot for a twinned network (red, pyramidal neurons; blue, interneurons), with weak internetwork (*I-E* and *E-I*) and strong intra-network (*II* and *E-E*) connections. *Bottom:* Time course of field potential (f.p.; simulated for pyramidal neurons at 250 μm from the network, tissue conductance of 100 mS cm^-1^; Methods) and [GABA]_e_, as indicated. Key model parameters: *N*=200 (100 pyramidal cells and 100 interneurons), *G*_ii_= 0.216 mS cm^-2^, *G*_ee_= 0.003 mS cm^-2^; Inter-network peak synaptic conductance *G*_ei_= 0.00012 mS cm^-2^ and *G*_ie_= 0.00064 mS cm^-2^, E-currents (Poisson series) *g*_si_=0.3 mS cm^-2^, tau = 3 ms, *f_s_*= 20 Hz; I-current (Poisson series) *g*_se_=0.003 mS cm^-2^, decay constant tau = 3 ms, *f*_s_= 20 Hz, *V_GABA_*=-58 mV, *A_f_* = 2×10^-7^ nS cm^-2^ ms^-1^, *G*_pump_=0.003 s^-1^.

We simulated networks of varied sizes and properties, including a plausible range of GABA release and uptake rates. First, simulations revealed that under constant *G*_tonic_, cells would fire stochastically, at a near-constant average frequency, showing little synchronization (Figure 3B). In striking contrast, when [GABA]_e_ (hence *G*_tonic_) dynamics were driven by interneuronal spiking, the network consistently exhibited spontaneous periodic activity patterns (Figure 3C-D), over a wide range of frequencies depending on several key parameters: the excitatory drive (*E*-current), synaptic connectivity strength (*G_ii_*), and the GABA release-uptake scaling factor *R* (Figure 3E; Methods). Simulations showed a robust, predictable relationship between interneuronal spiking, [GABA]_e_, and *G*_tonic_ during periodic busts of activity (Figure 3C-E, Figure S3). Interestingly, larger networks (containing 400-1500 cells) tended to have a more regular activity cycle (Figure S4), which is consistent with classical theory predicting less variability near a system’s Hopf bifurcation point (Fasoli et al., 2018), which occurs towards the peak of *G*_tonic_.

Finally, we explored the relationship between cell spiking activity, *G*_tonic_, and [GABA]_e_ in a twinned network configuration that incorporated both interneurons and principal cells (PCs) equipped with GABA_A_Rs. An initial exploration of connectivity parameters has indicated that such networks show stable behaviour under external stimuli when synaptic connectivity between the two networks (average synaptic conductance *G*_ie_, interneurons to PCs, and *G*_ei_, PCs to interneurons) were, on average, two orders of magnitude lower than intra-network connectivity (*G*_ee_ and *G*_ii_). Both interneuronal and PC networks were kept activated by stochastic excitatory conductance, *g*_si_ and *g_se_*, respectively. We found that, over a range of *g_se_* values, behaviour of the PC network was distinctly sensitive to [GABA]_e_-dependent *G*_tonic_ generated by the interneuronal network. PCs showed brief, highly synchronised bursts of activity that followed periodic rises of interneuronal spiking (Figure 3F). Strikingly, the PC spiking bursts occurred on the ascending phase of [GABA]_e_ waves (Figure 3F), thus displaying some distinct features of the epileptiform events that we observed experimentally (Figure 1D, Figure 2D).

### Self-maintained waves of tonic GABA_A_ conductance in hippocampal slices

To examine the validity of the causal relationships suggested by the modelling, we carried out several experiments. First, we asked whether the dynamics of *G*_tonic_ predicted by simulations (Figure 3A-D) occur in hippocampal circuits in the absence of glutamatergic transmission. We therefore recorded GABA_A_R-mediated currents from CA1 pyramidal neurons (whole-cell; unless specified otherwise, these and subsequent acute slice experiments were carried out in the presence of 50 μM NBQX, 10 μM APV, and 1 μM CGP52432, to block AMPA, NMDAR, and GABA_B_ receptors, respectively).

Elevating extracellular [K^+^] to 10 mM in nominally Mg^2+^-free medium, to increase interneuronal activity, elicited outward shifts in the holding current, which were sensitive to picrotoxin (a non-competitive GABA_A_R channel blocker; Figure 4A-B), and consisted of brief transients riding on periodic low frequency waves (Figure 4C). The frequency of the slow GABA_A_R-mediated waves (0.2-0.3 Hz) was similar to the periodic [GABA]_e_ transients observed with GABA imaging (Figure 2B). Consistent with the network model (Figure 3A-D, Figures S3-S4), the brief GABA_A_R-mediated currents, which most likely reflect individual interneuron discharges, tended to increase in intensity before the peak of the slow waves reflecting *G*_tonic_ (hence [GABA]_e_) (Figure 4C-E).

**Figure 4.**
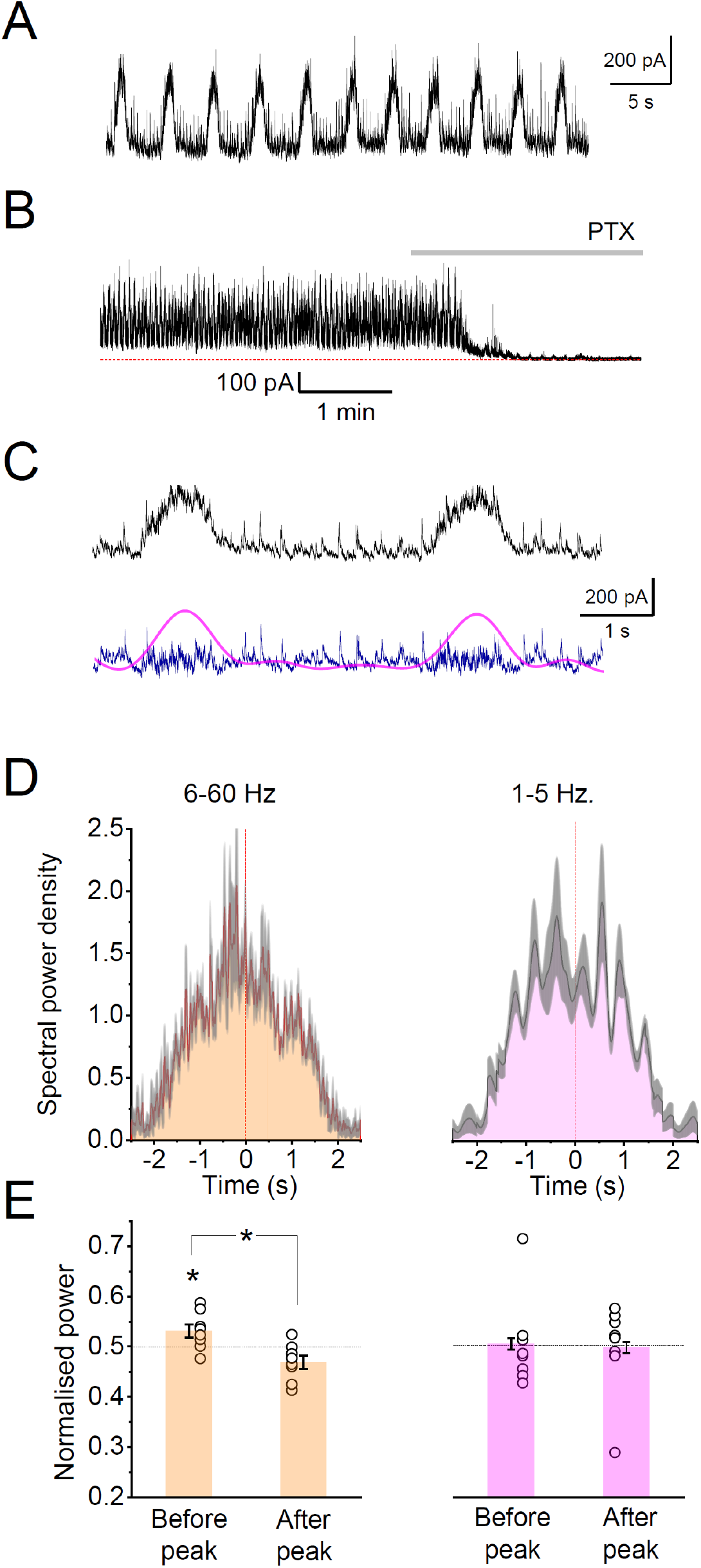
Experimentally induced slow wave-like oscillations of interneuronal activity. (**A**) Trace, example of GABA_A_R-mediated currents received by a CA1 pyramidal neuron (voltage-clamp with low intracellular Cl^-^; V_h_=0 mV). (**B**) Slow wave-like oscillations as in A are readily blocked by the GABA_A_R antagonist picrotoxin (PTX). (**C**) Traces as in A expanded: raw data (black) and its 2Hz low-pass (*G*_tonic_; magenta) and high-pass (blue) filtered components. (**D**) Example, average spectrogram plots (mean ± 95CI; high and low frequency bands, as indicated), data over multiple IIEs in one slice. (**E**) Summary of the analysis shown in D, for n = 7 slices: bar graphs, relative spectral power (bars, mean ± s.e.m.; dots, individual experiments), before and after the *G*_tonic_ peak, as indicated; *p < 0.05, one-sample (left) and paired-sample (middle) Student’s *t*-test.

### Synchronisation of fastspiking PV+ interneurons during GABA waves

To better understand the dynamics of synchronised interneuronal activity during GABA waves, we simultaneously monitored spiking activity in pairs of FS PV+ interneurons (in cell-attached mode), and GABA_A_R-mediated currents in a CA1 pyramidal cell (whole-cell mode), with ionotropic glutamate receptors blocked (Figure 5A). Clear rhythmicity of interneuron activity was documented in three out of nine experiments: wavelike activity (see Methods for definition) occurred in both cell-attached PV interneurons, *i.e*., 6 out of 18 recorded interneurons, suggesting that ~35% of PV+ interneurons could drive epileptiform discharges. This appeared in good correspondence with network simulations predicting that peak synchronisation involves up to 40% of interneurons (Figure 3C).

**Figure 5.**
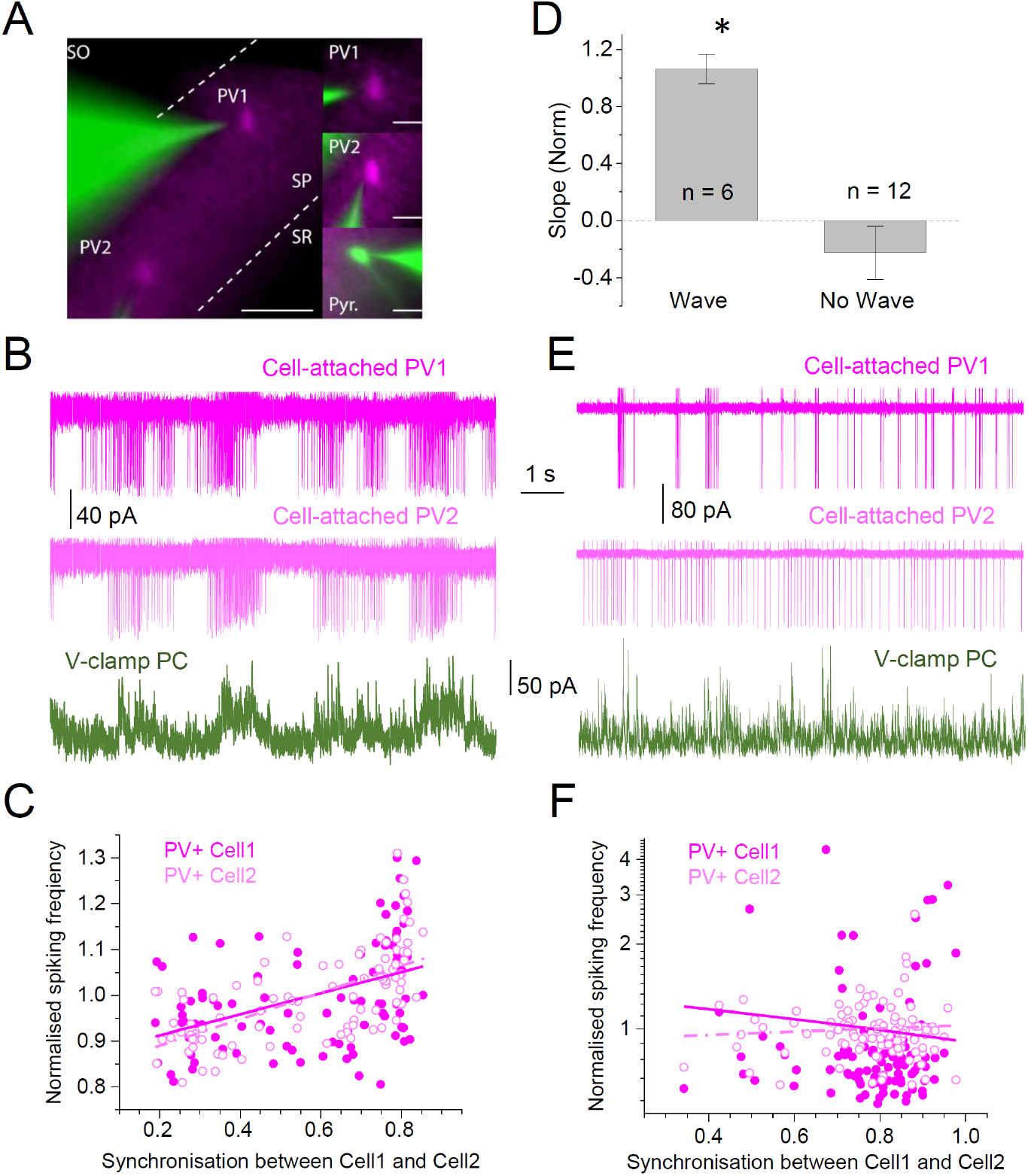
Synchronization of FS PV+ interneuron spiking activity during GABA waves. (**A**) Illustration of dual cell-attached recordings of PV+ interneurons (green: Alexa Fluor 488 50 μM; magenta: tdTomato) and voltage-clamp recording of a CA1 pyramidal cell (AF488, 50 μM). Scale bars, 50 μm (left) and 20 μm (right). (**B**) Example, time course of spiking activity of two simultaneously recorded PV+ cells (top and middle traces), and IPSCs in a pyramidal neuron (bottom, Vhold = +10 mV) in high K^+^ solution (10 mM) and 0 Mg^2+^; rhythmic network waves are present. (**C**) Spiking frequencies for the two PV+ cells (as shown in B), against their synchronization coefficient; straight lines, the corresponding linear regressions. (**D**) Summary, regression slope (mean ± s.e.m.) between PV+ spiking intensity and synchronization parameter in dual-patch recordings, as shown in C; samples with rhythmic waves (n = 6 cells in 3 slices) and without (n = 12 cells in 6 slices) are shown; *p < 0.03, unpaired Student’s t-test. (**E-F**) Examples as in (B-C) but with no detectable rhythmic waves; notations as in B-C.

In such experiments, quasi-periodic bursts of interneuron spiking (Figure 5B; magenta) were observed together with the corresponding fluctuations of GABAergic currents in the pyramidal neuron (Figure 5B, green). A systematic analysis of network activities showed a positive correlation between the spiking frequencies of the two PV+ cells and the synchronization parameter *k* (Methods; example in Figure 5C). Overall, experiments that featured prominent rhythmic waves also showed a significant interneuron synchronization-spiking correlation, whereas experiments that did not exhibit rhythmicity showed no such correlation (summary in Figure 5D; Figure 5E-F shows examples of experiments without rhythmicity, and related data).

Because highly synchronised firing of interneurons (as in Figure 5B) might suggest their coupling via gap junctions (Galarreta and Hestrin, 2001; Whittington and Traub, 2003), we asked if gap junctions could, at least in theory, prompt rhythm generation. In the network model containing 250 cells without GABA-wave signalling (as in Figure 3B), connecting neighbouring interneurons via gap junctions (individual conductance 0.1 mS/cm^2^) triggered network oscillations occurring at 10-100 Hz (Figure S5), but was unable to sustain the frequency range of 0.1-1 Hz characteristic of interictal activity, over the wide range of parameters illustrated above (Figure 3E).

### Spiking of principal neurons is synchronised shortly after the peak of interneuronal firing

Next, to examine how interneuronal network activity modulates firing of principal cells, we recorded one CA1 or CA3 pyramidal cell in cell-attached mode while another pyramidal neuron nearby was recorded in voltage clamp mode to monitor its GABAergic input (at 10 mM K^+^). Pyramidal cell spiking showed a characteristic lull during the rising phase of the *G*_tonic_ wave before rebounding as the latter reached a peak (Figure 6A left, blue segments). Blocking GABA_A_Rs with picrotoxin abolished this periodic fluctuation in spike frequency (Figure 6A right).

**Figure 6.**
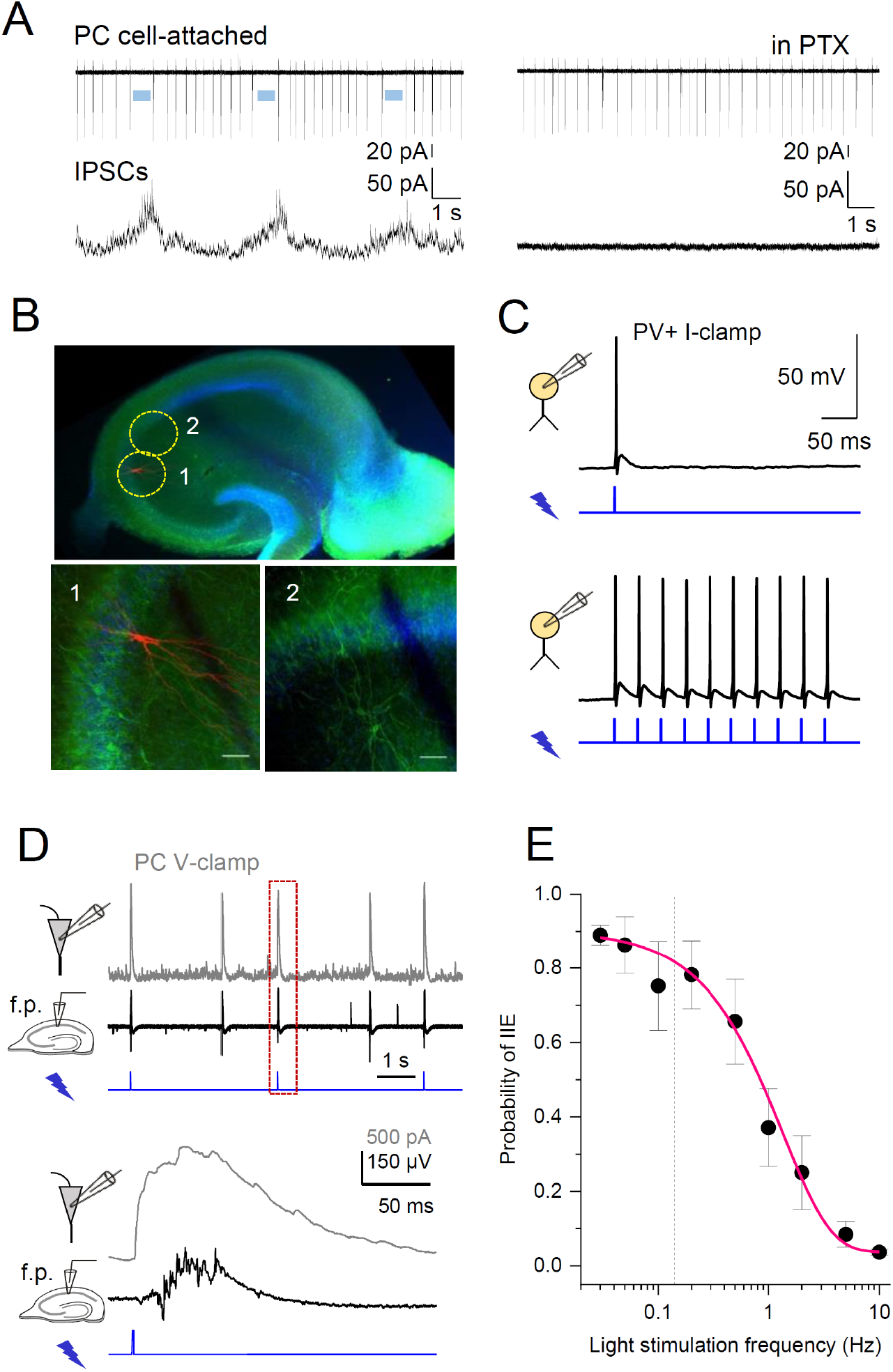
Short photoactivation of PV+ interneurons evokes epileptiform burst discharges. (**A**) Simultaneous cell-attached and whole-cell recordings of action potentials (top trace) and GABA_A_R-mediated IPSCs (bottom) during [GABA]_e_ waves (left) and after application of picrotoxin (PTX); blue segments indicate pauses in PC spiking before the peaks of IPSC. (**B**) Transverse hippocampal slice of PV:: *Cre* X Ai32 mouse(top): green fluorescence shows ChR2 expression in PV+ cells against DAPI (*blue*) nuclear counterstain (Methods). Images 1-2 (dotted circles above): *post hoc* identification of a CA3 pyramidal neuron (biocytin fill, *red;* magnified in 1) after a whole-cell recording shown below in D; scale bars: 20 μm. (**C**) Current-clamp recordings from a PV+ interneuron activated by a single (left) or repetitive (right) blue light pulses (1 ms, 470 nm). (**D**) Example of pyramidal neuron IPSCs (grey trace, top; Vhold = +10 mV) and field potentials (black trace, middle): photostimulation of PV+ cells (blue trace, bottom; 1 ms pulse) triggers large GABA_A_R currents and epileptiform bursts similar to those occurring spontaneously (e.g., Figure 1); 5 mM K^+^, 0-Mg^2+^ solution. (**E**) Statistical summary: Probability of evoking an interictal event (IIE, mean ± s.e.m., n = 3-7 slices for individual values, mean delay between the pulse and IIE: 43 ± 5 ms, 1 ms pulses) as a function of light stimulation frequency; the average frequency of spontaneous IIE (0.14 ± 0.03 Hz, n = 7 slices, dotted line) falls within the range of optimal frequencies to entrain the network with light stimuli; above 0.2 Hz stimulation, the network enters into a refractory state.

Thus, a rise in interneuron activity could paradoxically initiate interictal events in pyramidal cells, by synchronising their firing, consistent with the sequence of events reported earlier in the cortical circuitry (Librizzi et al., 2017). To test this further, we depolarised, synchronously and *en masse*, PV+ interneurons in the CA1 area using widefield optogenetic stimulation (Figure 6B). First, we confirmed that PV+ interneurons expressing channelrhodopsin-2 (ChR2-H134R-eYFP) reliably responded to 1 ms light pulses (Figure 6C). We next asked whether optogenetic stimulation of PV+ cells could prompt interictal events in the ‘epileptic’ slice (5 mM [K^+^], 0-Mg^2+^). Indeed, a single 1 ms light pulse evoked large GABA_A_R IPSCs in CA1 pyramidal neurons, with coincident epileptiform discharges (field potentials) that were indistinguishable from spontaneous interictal events (Figure 6D, Figure S6A). Again, this observation is consistent with that seen under similar circumstances, albeit a different experimental model of interictal activity, in cortical slices (Chang et al., 2018).

Blocking GABA_A_ receptors by bath application of picrotoxin completely abolished light-evoked interictal spikes, confirming a requirement for GABA release from PV+ cells (Figure 6 - figure supplement 1A). Notably, light stimuli also evoked spike bursts in pyramidal neurons, with a delay similar to that of interictal events (42 ± 3 ms *vs*. 43 ± 5 ms; Figure S6B). Thus, an abrupt release of PV+ cell inhibition upon light turnoff triggered synchronous pyramidal neuron discharges. Overall, 1 ms optogenetic stimuli applied at a frequency between 0.03 and 0.2 Hz consistently evoked neuronal bursts with ~0.8 probability whereas stimulation at > 0.2 Hz (value close to natural periodicity, 0.14 ± 0.03 Hz) was less effectual, with spike generation probability decreasing to ~0.1 at 5-10 Hz (Figure 6E; Figure S7).

### Evoked [GABA]_e_ waves can entrain network rhythms

We next asked whether an externally enforced wave of GABA release could entrain interictal discharges. To see if mimicking a relatively slow GABA wave, by a slow photoactivation of PV+ interneurons, could entrain the epileptic network, we applied ramp stimulation instead of brief square pulses. First, we carried out optogenetic excitation of PV+ cells using ramps (1-2 s duration, comparable to spontaneous GABA waves) while recording current activity in CA1 pyramidal cells, in control conditions (normal ACSF; Figure 7A). Ramp stimulation of PV+ cells elevated this activity only to a certain point, after which the IPSC frequency rapidly dropped (Figure 7B). In epileptogenic slices, similar events could be seen leading to interictal spikes (Figure 7C-D; Figure S8). Indeed, in five out of eight experiments, we clearly observed wave-like GABA_A_R-mediated currents induced by optogenetic ramp stimulation, with a probability of inducing interictal spikes with probability of 0.25 ± 0.09 and a delay of 831 ± 47 ms (detection window between 500 and 100 ms post-stimulation). In contrast, the three experiments without light evoked GABA waves exhibited a much lower likelihood of triggering epileptiform discharges (0.06 ± 0.04).#

**Figure 7.**
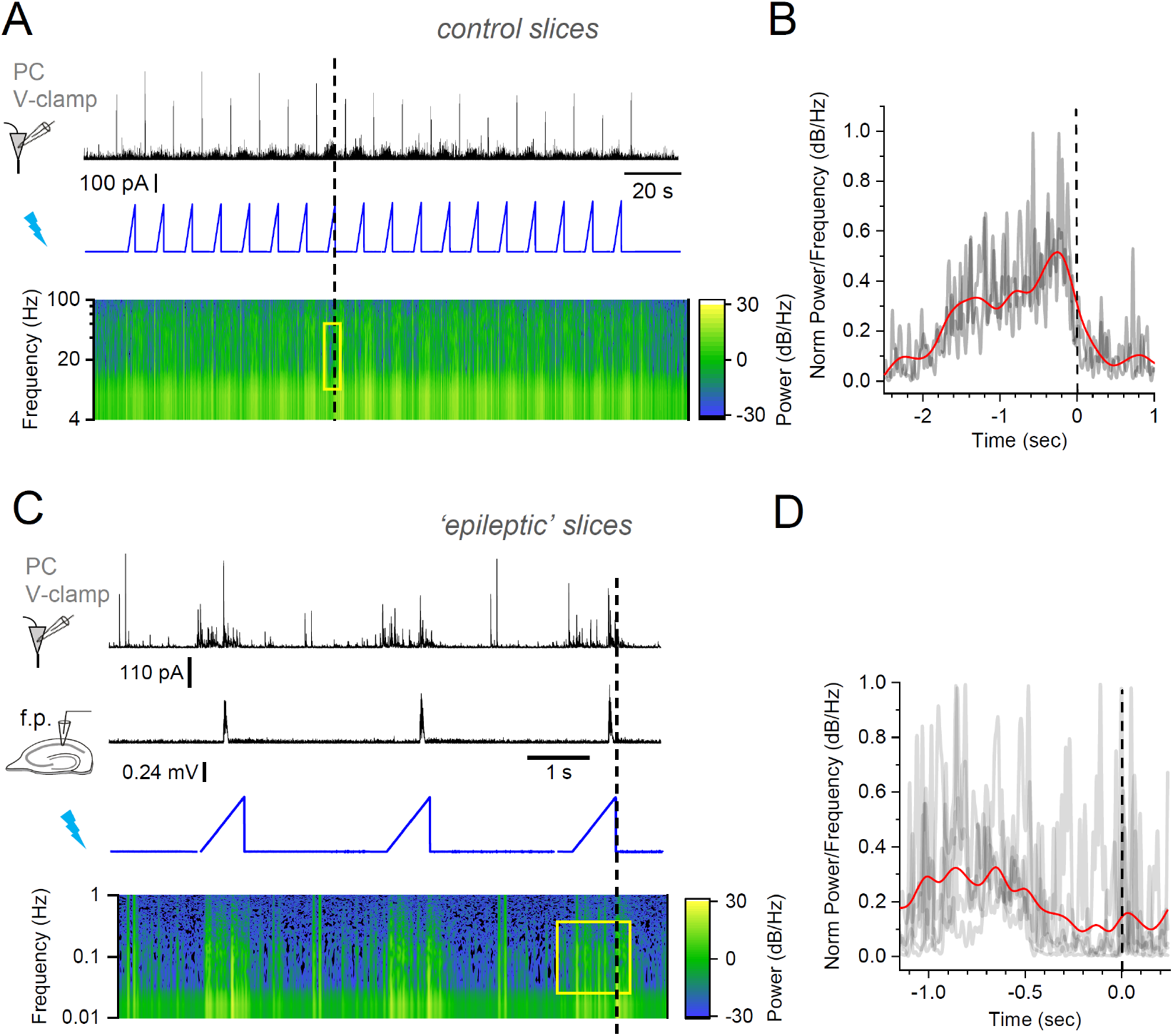
Optogenetically evoked GABA waves induce epileptiform burst discharges. (**A**) GABA_A_R current in pyramidal neurons (Vhold = +10 mV, normal aCSF solution; black trace, top) in response to progressive opto-activation of PV+ interneurons (blue, middle), with the IPSC power spectrum density (bottom); dotted line, end of individual light ramps. (**B**) Average spectral power density (5-50 Hz interval, n=18 ramp stimulations), time window as exemplified by yellow rectangle in A near light intensity peaks (dotted line). (**C**) Tests in epileptogenic tissue (0-Mg^2+^ and 5 mM K^+^ aCSF): GABA_A_R current in pyramidal neurons (black trace, top) and local field potentials (middle) during opto-activation (blue trace), with power spectrum density (bottom); note that interictal bursts occur towards the end of the stimulation ramp (vertical dotted line). (**D**) Average of spectral power density for time window exemplified by yellow rectangle in C; notations as in B.

To understand the biophysical basis of this phenomenon, we asked whether in our simulated network the *G*_tonic_ dynamics on its own could drive network synchronization and rhythmicity. We therefore forced *G*_tonic_ to follow a 1 Hz sine wave, and found that cell firing indeed peaked at *G*_tonic_ troughs, which were in turn preceded by maximal firing synchronicity (Figure S9), in line with the experimental data. Thus, increased interneuronal activity initially boosts network inhibition, followed by prevailing disinhibition (likely due to shunting), at which point synchronised network discharges are likely to occur, as generally predicted by our theoretical model.

### GABA uptake controls the frequency of periodic epileptiform events

The overall rate of GABA uptake depends on the numbers of available GABA transporters, dictating how rapidly [GABA]_e_ returns to its equilibrated steady-state level. The latter level, however, depends on the stoichiometry and kinetics of GABA transporters, not on their overall number. At the same time, changes in the uptake rate should have little effect on the rapid activation of intra-synaptic receptors (Rusakov, 2001). Thus, reducing the number of GABA transporters should decelerate the dynamics of [GABA]_e_ waves hence *G*_tonic_ without affecting synaptic GABAergic transmission. To test this principle theoretically, we explored a network of 1500 cells (as in Figure S3E-F) and found that increasing the rate of GABA uptake up to a certain level (~16 1/s) can drastically reduce the [GABA]_e_ dynamic range, thus effectively suppressing activity rhythms (Figure 8A-B, top panels). Conversely, slowing down GABA uptake allowed [GABA]_e_ to fluctuate rhythmically, with lower frequencies at lower decay rates (Figure 8A-B).

**Figure 8.**
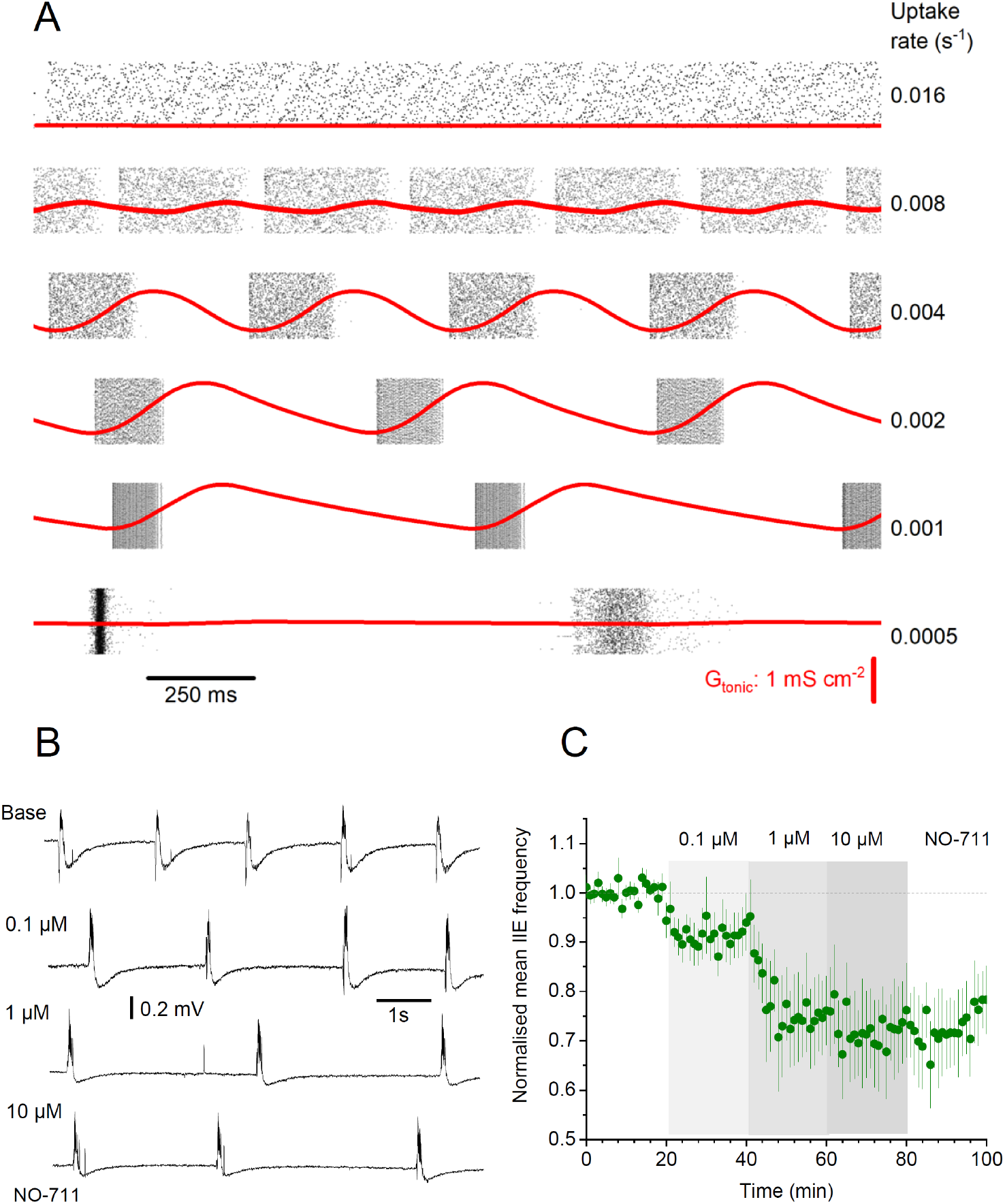
GABA uptake rate controls rhythmic activity of interneuronal networks. (**A**) Modelled time course of *G*_tonic_ at different GABA uptake rates, as indicated; key model parameters: cell number *N* = 1500, intra-network peak synaptic conductance *G*_ii_ = 0.096 nS cm^-2^; E-currents (Poisson series) with average synaptic conductance *g_s_* = 0.05 nS, decay constant tau = 3 ms, and frequency *f_s_* = 20 Hz; GABA_A_R reversal potential *V_GABA_* = −53 mV, GABA release factor *A_f_* = 10^-8^ nS cm^-2^ ms^-1^, *G*_pump_ =0.004 s^-1^ (see Methods for further detail); *G*_tonic_ scale (bottom panel) applies throughout. (**B**) Raster plots of interneuron spiking in the network, for the six cases shown in A; for presentation clarity, 6000 randomly selected spike events shown only, in each panel. (**C**) Examples of interictal spiking recorded in slice, under basal conditions (Base), and during subsequent bath applications of the GABA uptake blocker NO-711 at increasing concentrations, as indicated. (**D**) Average frequency of interictal events (mean ± s.e.m., n = 6 slices), normalised to the baseline value, in baseline conditions and during NO-711 application, as indicated.

To assess whether this causality is indeed present in real networks, we monitored interictal events in slices while adding a progressively increasing concentration of NO-711, a specific blocker of the main GABA transporter GAT-1, up to 10 μM, at which point its effect on [GABA]_e_ saturates (Kersante et al., 2013). This protocol was used to block increasing fractions of GAT-1, thus slowing down [GABA]_e_ reequilibration without affecting phasic GABAergic transmission or the basal (equilibrated ambient) [GABA]_e_ level. Strikingly, slowing down [GABA]_e_ uptake / reequilibration reduced the frequency of interictal discharges, in exact correspondence with the modelling predictions (Figure 8C-D). In some experiments, we also monitored local field potential and pyramidal neuron activity simultaneously, which again revealed rises in (tonic) GABA currents prior to field spikes (Figure S10). These results provide compelling evidence that [GABA]_e_ waves play a key role in the generation and pacing of network rhythms in interneuronal networks.

## DISCUSSION

We have found that interneuronal networks exhibit rhythmic synchronization at frequencies similar to interictal bursts, which also occurs in the absence of fast glutamatergic signalling. Cell excitation in interneuronal networks arises from the depolarising current through GABA_A_Rs, which depends on the relationship between receptor reversal potential and its synaptic activity-driven action (both phasic and tonic). This relationship involves the activity-dependent dynamics of intracellular chloride and extracellular potassium, thus engaging the key chloride extruder KCC2 and possibly other transporter mechanisms (Chavas and Marty, 2003; Song et al., 2011; Virtanen et al., 2021).

The exact nature of the underlying cellular machinery, which has been the subject of intense interest, was outside the scope of the present study. Here, we focused on the generation and pacing of network rhythms, proposing that it relies on periodic fluctuations of *G*_tonic_ driven mainly by activity-dependent dynamics of [GABA]_e_. Inhibitory restraint has earlier been proposed to “break down” due to depolarising block and reversal of the chloride potential. The present study identifies [GABA]_e_ fluctuations as an important mechanism underpinning initiation and propagation of ictal activity.

The waves of [GABA]_e_ provide a volume-transmitted feedback signal that modulates neuronal firing. It was earlier reported that reducing *G*_tonic_ from its basal (quiescent tissue) level boosts CA1 pyramidal cell firing while having little effect on interneuronal firing (Semyanov et al., 2003), consistent with ‘tonic’ disinhibition of principal cells. However, increasing *G*_tonic_ above a certain level could have a similar effect on principal cells as it shunts interneuronal activity (Song et al., 2011; Pavlov et al., 2014), at which point the disinhibition (rebound excitation) of excitatory neurons again prevails leading to synchronized network discharges. Interestingly, the earlier work (Pavlov et al., 2014) considered only a steady-state case in which [GABA]_e_ hence extra-synaptic GABA conductance was clamped, and no volume transmission of GABA occurred that would engage multiple cells. It was found that (a) low *G*_tonic_ had little effect on synchronization, (b) its initial increase boosted excitation and synchronization (mainly by damping excitatory noise; a high-pass filter effect), and (c) its further increase reduced excitability due to shunting, thus reducing synchronisation. Whilst that was a simple phenomenon isolated by having a stationary condition, the present work deals with the nonstationary network dynamics engaging [GABA]_e_ -dependent feedback: this leads to a different parameter space for synchronisation phenomena. Nonetheless, present simulations (Figure 3D, Figure 3 - figure supplement 1D) predict that synchronisation occurs in the rising phase of [GABA]_e_ ~ *G*_tonic_ while dropping near its peak, consistent with the earlier findings. Overall, the pattern of network activity and its predisposition to oscillations must depend on the dynamics of *G*_tonic_ driven by [GABA]_e_ release, diffusion, and uptake.

### [GABA]_e_ dynamic range and activation of extrasynaptic receptors

We find that during interictal discharges in hippocampal slices [GABA]_e_ peaks at about 1.5-2 μM. Calibration of the sniffer-patch sensor in acute slices indicates that [GABA]_e_ is maintained at 250-500 nM in nominally Mg^2+^-free aCSF. This concentration range is higher than [GABA]_e_ in quiescent slices (~100 nM), which we previously estimated using the same technique (Wlodarczyk et al., 2013), an observation possibly explained by a lower spontaneous firing rate of interneurons in the higher [Mg^2+^] concentrations. These estimates are generally in line with the [GABA]_e_ range reported earlier in hippocampal area CA1 (Farahmandfar et al., 2011). Are these levels of [GABA]_e_ compatible with peak values of *G*_tonic_ documented here using whole-cell recordings? One important prerequisite of having significant *G*_tonic_ is the presence of relatively low-affinity extrasynaptic GABARs that can stay in the sensitized state for prolonged periods of time. The well-described and broadly expressed high-affinity δ-subunit-containing GABA_A_Rs were found to become desensitized and almost irresponsive to GABA spillover when [GABA]_e_ reached ~0.25 μM (Bright et al., 2011). Other receptor sub-types with EC_50_ in the range of 0.5 - 2 μM include native metabotropic GABA_B_ receptors in L2/3 pyramidal neurons (Wang et al., 2010), and receptors with subunits α5β3γ2L (0.5-7 μM) and α1β2γ2L (0.6-9 μM) that can generate non-synaptic GABA currents in the hippocampus (Lagrange et al., 2018). In general, it therefore appears plausible that the [GABA]_e_ waves reported here find enough receptor targets to provide a significant dynamic range of *G*_tonic_ (Scimemi et al., 2005b).

### Mechanisms controlling [GABA]_e_

GABA that escapes from synaptic clefts (i.e., spills over) during interneuronal discharges is the most likely source of [GABA]_e_, although there have also been studies reporting GABA release from astroglia (Heja et al., 2009; Lee et al., 2011; Le Meur et al., 2012). One plausible mechanism which could regulate [GABA]_e_ is the reversal mode of GABA transporters. Four GABA transporter types (GAT1-3 and BGT1), all showing potassium- and voltage-dependent kinetics, are expressed in both neurons and glial cells (Borden, 1996; Conti et al., 1998; Gadea and Lopez-Colome, 2001). Intense excitatory activity, for instance during epileptiform bursts, can increase extracellular [K^+^] to 12-15 mM (Krnjevic et al., 1980; Somjen and Giacchino, 1985) and could, in principle, reverse GABA transport, thus generating GABA efflux (Wu et al., 2001). As this type of condition is compatible with the present experimental settings, GABA transporters could potentially be a source of extracellular GABA coming from both neurons and glia. However, any significant rises of [GABA]_e_ are likely to prevent the reversal transporter mode (Savtchenko et al., 2015b). This argues against a significant contribution of non-synaptic, transporter-mediated GABA release in conditions of our experiments. Finally, regulation of [GABA]_e_ could arise from the Bestrophin 1 Ca^2+^-dependent Cl^-^ channel (Lee et al., 2010) found predominantly in glial cells (Oh and Lee, 2017). However, this mechanism is unlikely to play a role here, since the operation of this channel does not appear correlated with neuronal activity (Lee et al., 2010).

### Constraining parameters of simulated neural networks

Our study explored a well-established FS interneuron network model (Wang and Buzsaki, 1996) that incorporates *G*_tonic_ driven by activity-dependent [GABA]_e_ kinetics (Pavlov et al., 2014; Savtchenko and Rusakov, 2014). Whilst this model replicated key aspects of the experimentally documented interictal activity in the hippocampus, it had certain limitations. Firstly, we used a near-linear relationship between [GABA]_e_ and *G*_tonic_, whereas in reality it incorporates a more complex, supra-linear dependence (Song et al., 2011; Pavlov et al., 2014). This might explain why the simulated *G*_tonic_ dynamics appears more wave-like than the experimentally observed regular [GABA]_e_ rises. Secondly, in the majority of simulations we implemented a simple stochastic excitatory input to mimic synaptic activity of pyramidal neurons, which is thus set independent of interneuronal activity. This approach was taken partly because the main aim of the present study was to establish whether and how the network activity of FS interneurons, on its own, could drive hippocampal oscillations. Nonetheless, a twinned simulated network containing both interneurons and principal cells displayed spiking behaviours that were fully consistent with the key features of interneuronal networks, and, more importantly, with experimental observations.

Finally, astroglia have also been considered a potentially important source of extracellular GABA due to reversal-mode release through GAT-1 and GAT-3 transporters. However, as pointed out above, such release can only occur at the lowest levels of [GABA]_e_, hence out-of-phase with the interneuron activity, whereas our experimental observations combined indicate the opposite trend. Nonetheless, the role of astroglial GABA uptake must play an important role in pacing network rhythms, as suggested by both simulations and experiments that established a clear relationship between the GABA uptake rate and the frequency of epileptiform events.

### Interneurons and principal neurons in rhythm initiation

Our sniffer-patch experiments, GABA imaging, and recordings of FS PV+ interneuron activity point to an increased tonic inhibition before interictal discharge onset. This observation invokes two possibilities: either interictal activity is due to the disinhibition of principal cells (due to the ‘self-inhibition’ of the interneuronal network or to depolarisation block) or interneuron firing itself drives interictal discharges. Our experiments reveal that synchronised interneuronal network activity can determine the timing of co-ordinated pyramidal cell bursts, confirming the earlier suggestion that interneurons provide local spatiotemporal control of pyramidal neurons and are critically important for generating such network rhythms (Cobb et al., 1995; Traub et al., 1999). At the same time, interneuronal activity *per se* can make an important contribution to epileptiform discharges.

Different areas of the brain display distinct combinations of physiological processes involving oscillations of interneuron networks. We would like to highlight two characteristic mechanisms behind such periodic activities, which may or may not depend on [GABA]_e_ and *G*_tonic_. Firstly, the so-called giant depolarising potentials (GDPs), which are network-driven wave-like events occurring during development (Sipila et al., 2005; Sipila et al., 2006; Khalilov et al., 2015). The average GDP duration is around one second, with a fairly random spiking pattern involving 6-10 discharges per minute (Leinekugel et al., 1998; Kasyanov et al., 2004). It has been suggested that GDPs are generated by synergistic activity of pyramidal cells and interneurons (Ben-Ari et al., 2007). The second mechanism involves high-frequency (up to 200 Hz) oscillations, referred to as hippocampal ripples (Hulse et al., 2016; Oliva et al., 2016; Roux et al., 2017), and represent highly synchronous bursts of brain activity. In acute slices of ventral hippocampus, such ripples can occur in a periodic pattern, with a frequency of 7-14 Hz (Papatheodoropoulos, 2010), similar to the theta-rhythm. These oscillatory patterns can thus arise purely from the interaction of excitatory and inhibitory neurons, a mechanism qualitatively different from interictal activity resulting from periodic fluctuations in [GABA]_e_ and *G*_tonic_.

## ACKNOWLEDGEMENTS

This work was supported by Epilepsy Research UK, MRC, Wellcome Trust Principal Fellowship, ERC Advanced Grant. Optimization and parallelization of ARACHNE algorithms for extended neural network simulations was provided by Andrey Galkin and team at AMC Bridge (Waltham, MA).

## AUTHOR CONTRIBUTIONS

I.P., L.P.S. and D.A.R. narrated the study; I.P. and V.M. designed and carried out electrophysiological and optogenetic studies; N.C. carried out iGABASnFR imaging experiments designed and analysed by T.P.J; S.S. carried out sniffer-patch experiments; L.P.S. designed and carried out network modelling studies and data analyses; J.S.M. and L.L.L. developed iGABASnFR imaging protocols; D.M.K. and M.C.W. designed optogenetic experiments and assisted with analyses; D.A.R. designed selected experiments and simulations and wrote the manuscript with I.P. and L.P.S., which was subsequently contributed to by V.M. and all other authors.

## DECLARATION OF INTERESTS

The authors declare no competing interests

## METHODS

### Animal experimentation

All experiments involving animals were carried out in accordance with the European Commission Directive (86/609/EEC) and the United Kingdom Home Office (Scientific Procedures) Act (1986), under the Home Office Project Licence PPL P2E0141 E1.

### Acute slice preparation

*In vitro* electrophysiological recordings were performed in acute hippocampal slices prepared from 3- to 4-week-old male Sprague-Dawley rats (Harlan Laboratories Inc, Bicester, UK). Animals were kept under standard housing conditions with 12 h light-dark cycle and free access to food pellets and drinking water. After being sacrificed using an overdose of isoflurane, animals were decapitated, brains were rapidly removed, and hippocampi were dissected for slice preparation. Transverse hippocampal slices (350 μm-thick) were cut with a Leica VT1200S vibratome (Germany) in an ice-cold sucrose-based solution containing (in mM): sucrose (70), NaCl (80), KCl (2.5), MgCl_2_ (7), CaCl_2_ (0.5), NaHCO_3_ (25), NaH_2_PO_4_ (1.25), glucose (22), bubbled continuously with 95% O_2_ + 5% CO_2_ to yield a pH of 7.4. Slices were allowed to recover in a sucrose-free artificial cerebrospinal fluid (aCSF) solution (in mM): NaCl (119), KCl (2.5), MgSO_4_ (1.3), CaCl_2_ (2.5), NaHCO_3_ (26.2), NaH_2_PO_4_ (1), glucose (22), bubbled with 95% O_2_ and 5% CO_2_ in an interface chamber for at least 1 hour at room temperature before being transferred to a submerged recording chamber.

Modified aCSF (nominally 0 mM Mg^2+^ and 5 mM K^+^, unless specified otherwise) was used to induce epileptiform activity. To facilitate the rapid generation of epileptiform discharges slices were perfused with the solution on both sides. All recordings were done at 32°C. GABA_B_ receptors were blocked in all experiments by 5 μM CGP52432. Field potential recordings from stratum pyramidale were performed with 1-2 MΩ glass electrodes filled with aCSF. Visualized whole-cell voltage-clamp recordings were performed from CA1 pyramidal neurons using an infrared differential interference contrast imaging system. Recording pipettes (3-5 MΩ) were filled with internal solution, containing (in mM): Cs-methanesulfonate (120), HEPES (10), EGTA (0.2), NaCl (8), MgCl_2_ (0.2), Mg-ATP (2), Na-GTP (0.3), QX-314 Br-salt (5), pH 7.2, osmolarity 290 mOsm kg^-1^. GABA_A_R-mediated currents were recorded from neurons voltage clamped at 0 mV or +10 mV (close to the reversal potential of glutamatergic currents). To further prevent the contribution of NMDA receptors MK801 (1 μM) was included in the recording pipettes in acute slices experiments. Areas CA3 were cut off in the outside-out patch experiments, but not in other settings: this had no detectable effect on interneuronal network activity in area CA1.

Series resistance (Rs) was monitored throughout the experiment using a −5 mV step command. Cells showing a >20% change in Rs, or values >25 MΩ, or an unstable holding current, were rejected. Recordings were obtained using a MultiClamp 700B amplifier (Molecular Devices, CA, USA), filtered at 4 kHz, digitized and sampled through an AD converter Digidata 1550 (Molecular Devices) or NI PCI-6221M (National Instruments) at 10 kHz and stored on a PC. Data acquisition and off-line analysis were performed using WinEDR 3.0.1 (University of Strathclyde, Glasgow, UK) and Clampfit 10.0 (Molecular Devices Corporation, USA) software, or Femtonics MES (Femtonics, Budapest) and custom-made Matlab (Mathworks) scripts running in Matlab r2020b.

### Outside-out patch recordings

Single channel GABA_A_R-mediated currents were recorded in the presence of 0.1 μM CGP-55845 as detailed earlier (Wlodarczyk et al., 2013; Sylantyev et al., 2020). Outside-out patches were pulled from dentate granule cells, raised above the slice surface and moved to area CA1; recordings were performed in voltage-clamp mode at 33–35°C. For membrane patches held at −70 mV, the recording electrode solution contained (in mM): 120.5 CsCl, 10 KOH-HEPES, 10 BAPTA, 8 NaCl, 5 QX-314-Br^-^ salt, 2 Mg-ATP, 0.3 Na-GTP; pH 7.2, 295 mOsm kg^-1^. For membrane patches held at 0 mV, the recording electrode solution contained (in mM): 120 Cs-methanesulfonate, 10 HEPES, 0.2 EGTA, 8 NaCl, 0.2 MgCl_2_, 2 Mg-ATP, 0.3 Na-GTP, 5 QX-314-Br^-^ salt; pH 7.2, 290 mOsm kg^-1^. Recordings were made using MultiClamp 700B amplifier (Molecular Devices, CA, USA); signals were digitized at 10 kHz. The patch pipette resistance was 5–7 MΩ. In experiments where GABA EC50 was determined, we used a θ-glass application pipette with ~200 μm tip diameter attached to a micromanipulator. The pipette position was controlled by a piezoelectric element (each switch took 50–100 μs) to allow rapid solution exchange. One pipette channel was filled with the bath aCSF solution; another channel had aCSF with varying concentrations of GABA. The flow rate was driven by gravity (Sylantyev and Rusakov, 2013b).

### Analysis of single-channel recordings

The frequency of GABA_A_R channel openings was calculated as *N /Δt*, where *N* is the number of openings and Δt is the time of recording. *N* was counted using a detection threshold of 1.5 pA above the mean baseline value and a minimum opening time of >0.2 ms. Single channel conductance was calculated as *G* = *I*/ (*V_rev_ - V_hold_*), where *I* stands for receptor-mediated current, and *V_rev_* – calculated chloride reversal potential. The average charge transfer through receptors was calculated as *Q* = *G*× *N* / Δt. The concentration-effect plot was fitted with the Hill equation 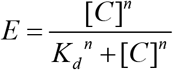, where *E* stands for the percentage of maximal charge transfer, *C* concentration of GABA, *K_d_* microscopic dissociation constant (equal to EC_50_), and *n* Hill’s coefficient.

### Organotypic slice culture preparation and biolistic transfection of iGABASnFR

Organotypic hippocampal slice cultures were prepared and grown with modifications to the interface culture method from P6–8 Sprague-Dawley rats as previously described (Jensen et al., 2019). In brief, 300 μm thick, isolated hippocampal brain slices were sectioned using a Leica VT1200S vibratome in ice-cold sterile slicing solution consisting of (in mM) Sucrose 105, NaCl 50, KCl 2.5, NaH_2_PO_4_ 1.25, MgCl_2_ 7, CaCl_2_ 0.5, Ascorbic acid 1.3, Sodium pyruvate 3, NaHCO_3_ 26 and Glucose 10. Following washes in culture media consisting of 50% Minimal Essential Media, 25% Horse Serum, 25% Hanks Balanced Salt solution, 0.5% L-Glutamine, 28 mM Glucose and the antibiotics penicillin (100U/ml) and streptomycin (100 μg/ml), three to four slices were transferred onto each 0.4 μm pore membrane insert (Millicell-CM, Millipore, UK). Cultures were then maintained at 37°C in 5% CO_2_ and fed by medium exchange every 2-3 days for a maximum of 21 days in vitro (DIV). At 5DIV cultures were treated overnight with 5 μM cytosine arabinoside (Ara-C, a poison inhibitor of DNA and RNA polymerases) to reduce glial reaction following biolistic transfection and returned to standard culture media at 6DIV. At 8DIV cultures were shot with 1.6 micron Gold micro-carriers coated with 30 μg of hSyn.iGABASnFR.F102G plasmid using the Helios gene-gun system (Bio-Rad). Following transfection cultures remained for 5-10 days before experiments were carried out.

### Imaging of interictal epileptiform activity associated extracellular GABA transients

For imaging of iGABASnFR transients associated with interictal activity, organotypic slice cultures were cut from their membrane insert and transferred to the stage of a Femtonics Femto3D-RC imaging system (Femtonics, Budapest) integrated with patch-clamp electrophysiology and linked on the same light path to two femtosecond pulse lasers MaiTai (SpectraPhysics-Newport) with independent shutter and intensity control as previously described (Jensen et al., 2019). Slices were continuously perfused with bicarbonate based artificial cerebrospinal fluid (aCSF) equilibrated with 95% O_2_ and 5% CO_2_ at 32–34 °C using a gravity driven perfusion system (flow rate 3-4ml/min). aCSF solution contained (in mM): 120 NaCl, 10 glucose, 2.5 KCl, 1.3 MgSO_4_, 1 NaH_2_PO_4_, 25 NaHCO_3_, 2 CaCl_2_ with osmolality of 300 ± 5 mOsm kg^-1^. Biolistic transfection of iGABASnFR resulted in sparse labelling of neurones in the CA1 and CA3 regions making identification of iGABASnFR expressing cells using two-photon imaging (λ_×_^2p^ = 910 nm) unambiguous. Following identification, a glass field recording electrode filled with standard aCSF was placed within ~30 μm of the cell of interest and interictal activity was then initiated by exchanging the extracellular solution for a 0 mM Mg^2+^ aCSF (otherwise the same composition). The local field potential was then monitored until stable interictal bursting was observed, at which point curved frame-scan regions of interest following the somatic membrane were chosen for scanning (see Figure 2A for illustration). To avoid photodamage, scan duration was limited to 9 second epochs which were repeated with a one-minute intervals until a minimum of 10 events were captured. For comparison of interictal event timing and associated iGABASnFR transients; interictal event peak times were first identified by the findpeaks function in MATLAB and were accepted for further analysis where single peaks with a minimum width of 20ms and amplitude >5 x standard deviation (SD) of baseline noise could be detected. The associated iGABASnFR ΔF/F peak was then identified (again using *findpeaks*) within a manually adjusted window −750 ms to + 2s (extremes between cells/slices) over the peak times identified for the interictal event, with a minimum >2.5 x SD baseline and minimum width of 50ms. Events within that window were then analysed where a clear and stable iGABASnFR *ΔF/F* 50ms length baseline, rise and visible decay associated could be observed. The rise to peak from baseline was then fitted by polynomial regression and the 20% iGABASnFR peak time extrapolated from the fit and used to calculate iGABASnFR transient-interictal event lag times.

### Targeted cell-attached recordings and optogenetic experiments

PV::cre mice (B6;129P2-Pvalb^tm1(cre)Arbr^/J Jackson laboratory stock number: 008069) were crossed with Ai9 or Ai32 mouse line, which has floxed-stop tdTomato (B6.Cg-Gt(ROSA)26Sortm9(CAG-tdTomato)Hze/J Jackson laboratory stock number: 007909) or EYFP-tagged excitatory opsin channelrhodopsin-2-H134R (B6;129S-Gt(ROSA)26Sor^tm32(CAG-COP4*H134R/EYFP)Hze^/J Jackson laboratory stock number: 012569), to produce animals expressing tdTomato or channelrhodopsin-2 (ChR2) in parvalbumin-positive (PV^+^) interneurons throughout the brain (Madisen et al., 2012). Animals were kept under standard housing conditions with 12 h light-dark cycle and free access to food pellets and drinking water. Hippocampal slices were prepared from mice of both sexes aged between postnatal day 25 and 50 for the electrophysiology section, but under low light intensity to minimise any adverse effect due to activation of ChR2. Field potential recordings from stratum pyramidale and visualized whole-cell voltage-clamp recordings were performed from CA1 pyramidal neurons using infrared differential interference contrast imaging system as in the electrophysiology section. For the dual cell-attached recordings (Figure 5), borosilicate glass pipettes (3-5 MΩ) filled with aCSF and containing Alexa Fluor 488 (50#μM) were used and the whole-cell recording pipettes was containing the standard Cs-methylsulfonate cited above with the addition of Alexa Fluor 488. In addition, the slices were superfused with aCSF with 0 mM Mg^2+^ and 10 mM K^+^ as well as APV (50 μM), NBQX (10#μM) and CGP52432 (1 μM). #

Cells showing a series resistance >25 MΩ, or an unstable holding current, were rejected. Recordings were obtained using a MultiClamp 700B amplifier (Molecular Devices, CA, USA), filtered at 2-4 kHz and digitized at 10 kHz. Data acquisition and off-line analysis were performed using WinEDR 3.0.1 (University of Strathclyde, Glasgow, UK) and Clampfit 10.0 (Molecular Devices Corporation, USA) softwares as well as MATLAB and Python custom-scripts.

Wide-field illumination of the CA1 region of the hippocampus was delivered through a 20x water immersion objective (Olympus). PV tdTomato-positive cells were visualised with 590 nm wavelength and ChR2 was activated by blue light (wavelength 470 nm) generated by pE-2 LED illumination system (CoolLED); light intensity under the objective was in the range of 5 - 10 mW. For the ramp stimulation protocol, a train of 1 s ramps was delivered at on average ~0.3 Hz ranging from 0.12 to 0.33 Hz.

In some experiments (Figure 5), neurons showing the ‘wave-like’ activity were defined as follows. The voltage clamp trace was lowpass filtered with an IIR Butterworth filter at a cutoff frequency of 2Hz. The FFT was then computed on the filtered trace and if a significant (above 95CI) peak was present on the power spectrum between 0.01-2Hz, the activity was considered as wave-like.

### Interictal discharge analysis

The detection of interictal spikes was done using custom-made python script. Local field electrophysiological trace was first downsampled to 1 kHz and filtered using a Butterworth bandpass filter from 1 to 50 Hz. The onset of interictal discharges was determined by using a detection threshold on the second derivative of the local field potential recording. The threshold corresponded to a minimum of 4 SDs above baseline noise (taking from the beginning of each trace). For the probability of inducing an interictal events with 1 ms light pulse stimulation, a window of 100 ms post-stimulation was considered for the detection of an event. Concerning the ramp stimulation, the window was between 500 ms after the start of the ramp and 100 ms post-stimulation.

### Modelling: Interneurons and pyramidal neurons networks

The interneurons and pyramidal neuron (principal cell, PC) networks were simulated on a digital platform ARACHNE with remotely-controlled parallel computing (Aleksin et al., 2017). Similar to the previously explored non-hierarchical networks (Pavlov et al., 2014), the present one featured circular connectivity (Figure 3A), which helped (a) exclude edge effects, (b) equalise cell contribution, and (c) represent the size by a single parameter, network radius *R*. In individual simulated cells, the channel kinetics were typical of hippocampal fast-spiking basket interneurons (Wang and Buzsaki, 1996). Other incorporated biophysical mechanisms, such as ion channel flux, GABA transporter, synaptic and GABA tonic current were in accord with experimental findings (Gloveli et al., 2005; Tort et al., 2007; Kopell et al., 2010), as detailed below. The time course of GABAergic synaptic conductance followed the function *y*(*t*) = *G_ii_* (exp(-*t*/*τ_1_*) – exp(–*t*/*τ*_2_)) where *y*(*t*) is the synaptic conductance at time *t*, *τ*_1_ is the rise time constant, *τ*_2_ (termed tau elsewhere) is the decay time constant, *G_ii_* is the peak conductivity (mS cm^-2^) of synapses.

A network of interneurons and pyramidal cells was simulated using previously published models (Tort et al., 2007; Kopell et al., 2010), which was further developed as a full-scale cloud-computing platform ARACHNE (Aleksin et al., 2017), with minor modifications of the internal parameters. The critical addition was the dynamic *G*_tonic_, as explained below, with the *G*_tonic_ range in interneurons twice that of pyramidal cells, in accord with experimental data (Pavlov et al., 2014); the basic cell-circuit model used in the present study was obtained from ModelDB (accession no. 138421). Briefly, the modelled circular network consisted of *N* fast-spiking interneurons (*I*-cells) and *M* pyramidal cells (*E*-cells). Each cell was modelled as a single compartment using standard Hodgkin–Huxley formalism. The GABA_A_R reversal potential *V_GABA_* was set between −55 mV and −72 mV, as specified. Unless specified otherwise, the network featured three connection types: *I–I, I–E*, and *E–I*, with release probabilities values *P_E–E_* = 0.5, *P_E–I_* = 0.5, and *P_I–I_* = 1.0, respectively. The spatial synaptic connection density was defined as the matrix *W_XY_*(*i,j*) with the Gaussian distribution centred at a given neuron, so that

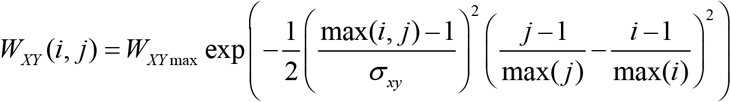

The model consists in bell-shaped strength and Gaussian density of connections. Synaptic conductance matrices of this model are with following strength coefficient *W_ee,max_* = 2, *W_ii,max_* = 0.8, *W_ei,max_* = 0.9; *W_ie,max_* = 0.3, and spatial SD: σ_ee_ = 10 μm, σ_ei_ = 12.5 μm, σ_ie_ = 8 μm and σ_ii_ =5 μm. The size (radius) of interneuron and principal cell network was set at 250 and 200 μm, respectively, with the signal propagation speed at 0.1 μm ms^-1^.

Simulations were performed using ARACHNE on the Amazon AWS cloud computing (cluster c5.large, tolerance 10^-5^, time step 0.02 ms). Random generator *use32BitRng* (MATLAB) was set to generate a delta correlated white noise for any stochastic processes. The initial voltage of interneurons in the network was set uniformly randomly, between −73 and −67 mV. Mechanisms of synaptic plasticity were excluded from the basic model of the neural network.

### Modelling: Network synchronization

The network synchronization parameter *k*(*τ*), which was also used for the experiment in PV+ cell pairs (Figure 5), was calculated as an average of all coefficients *k_ij_*(*τ*) for each pair of neurons (*i,j*). The time window of synchronization was divided into bins so that *τ* = 0.1/*f_m_* where *f_m_* is the network-average spiking frequency (see below). In each bin, an action potential was represented in a binary format (yes-no series). Next, *k_ij_*(*τ*) was calculated for each pair of neurons for the time bin *τ* using the formula:

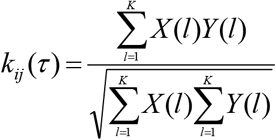

where *X*and *Y* are the binary series of the *i*th and *j*th cells, respectively, *l* is the bin number; thus, *X*(*l*) and *Y*(*l*) are either 0 or 1 depending on having a spike (1) or no spike (0) event for *i*th or *j*th during the *l*th bin, and *K* is the total number of bins.

### Modelling: Equations for the tonic GABA conductance

[GABA]_e_ activates extrasynaptic GABA_A_Rs, which generate *G*_tonic_. Thus, modelled cells generate non-specific tonic current: *I_tonic_* = *G_tonic_*(*V_GABA_ – V_m_*), where *V*_GABA_ GABA_A_R reversal potential, and *V_m_* is the membrane potential. *G_tonic_* depends on the interneuron network firing frequency, which in turn generates [GABA]_e_ increments, in accord with:

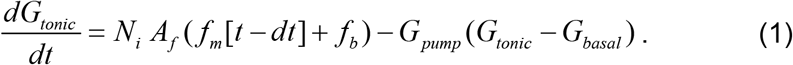

with initial *G_tonic_*(0) = *G_basal_* and *f_m_* is calculated as the total of cell-generated spikes over time *T:*

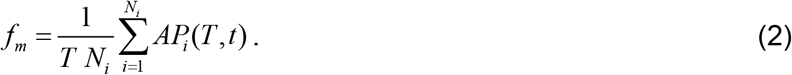

Other parameters were: *N*(or *N_i_*), total number of neurons in the network; *A_f_* (0.01 nS cm^-2^ ms^-1^, unless specified otherwise), tonic GABA conductance resulting from a single AP over 100 ms; *G*_pump_ (values as indicated), the GABA uptake rate, *G*_basal_ = 0.1 mS cm^-2^, [GABA]_e_ value at which GABA transporters reverse, *f_b_* = 0.1 Hz, basal average network spiking frequency at rest; and *dt*, a delay between neurotransmitter release and activation of extrasynaptic receptors (*dt* was <1 ms throughout).

In our simulations, we assumed that *G_tonic_* linearly depends on [GABA]_e_: *G*_tonic_ = α[GABA]_e_. This assumption provides a good approximation [GABA]_e_ < 1-2 μM, as in this case it falls into the near-linear approximation of the charge transfer versus [GABA]_e_ relationship (Figure 1E). The *G*_tonic_ dynamics was therefore determined by the time course of [GABA]_e_, which is represented by quasi-instantaneous release followed by uptake (i.e., quasi-exponential decay of [GABA]_e_ to its basal level). The latter was set in accord with previous estimates of the GAT-1 kinetics (Savtchenko et al., 2015b), so that solution of equation (1) for constant *f_m_* was

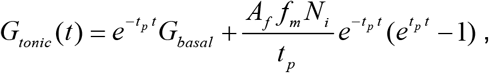

where *t_p_* is 0.02-0.1 ms^-1^ and *A_f_* is the *G*_basal_ / *G*_tonic_ ratio at steady-state for a given frequency 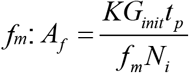, where 2 < *K* < 4. Thus, *A_f_* was a scaling factor for [GABA]_e_ action.

### Modelling: [GABA]_e_ dynamics

The dynamics of [GABA]_e_ was estimated from the cell-spiking raster plot, with individual action potentials releasing GABA at a rate of 0.5 μM ms’^1^ of GABA (~3000 molecules) in an extracellular volume of 20 μm radius, the average distance between neighbouring neurons of the network. [GABA]_e_ decays due to GABA uptake and diffusion, a typical constant of 0.004 μM ms^-1^ (ref. (Savtchenko et al., 2015b)), unless specified otherwise.

To explore the parameter space with respect to rhythm generation, and for the sake of clarity, we introduced a single dimensionless parameter *R* as a combined GABA release-uptake factor incorporating *A_f_* and *G_pump_*, in accord with

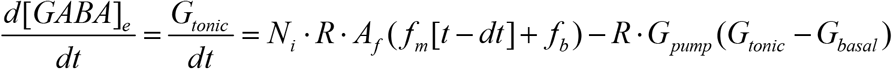

were term notation is as described above.

### Modelling: Code availability

ARACHNE is available with explanatory documentation at www.neuroalgebra.com. The program is made available under MIT license. ARACHNE is written in a way that allows users to run it on any remote platforms. Files with a set of initial parameters for reproducing the results compatible with ARACHNE can be provided upon request.

For any computations we provided the .exe file of ARACHNE, but also the initial code on https://github.com/LeonidSavtchenko/Arachne.

### Statistics

Shapiro-Wilk tests for normality were routinely run for small samples (this test for the means could be misleading for n > 15-19 due to Central Limit Theorem). Correspondingly, two-tailed paired and unpaired Student’s *t*-test, or otherwise non-parametric Mann-Whitney tests were used for statistical analyses. Mean difference was considered significant at the null-hypothesis rejection level of *p* < 0.05. Statistical summary data are shown as mean ± s.e.m. unless specified otherwise.

## Notes

### Competing Interest Statement

The authors have declared no competing interest.

### Summary of Updates

Text revisions

