## Supplementary Figures for "Volume-transmitted GABA waves pace epileptiform rhythms in the hippocampal network"

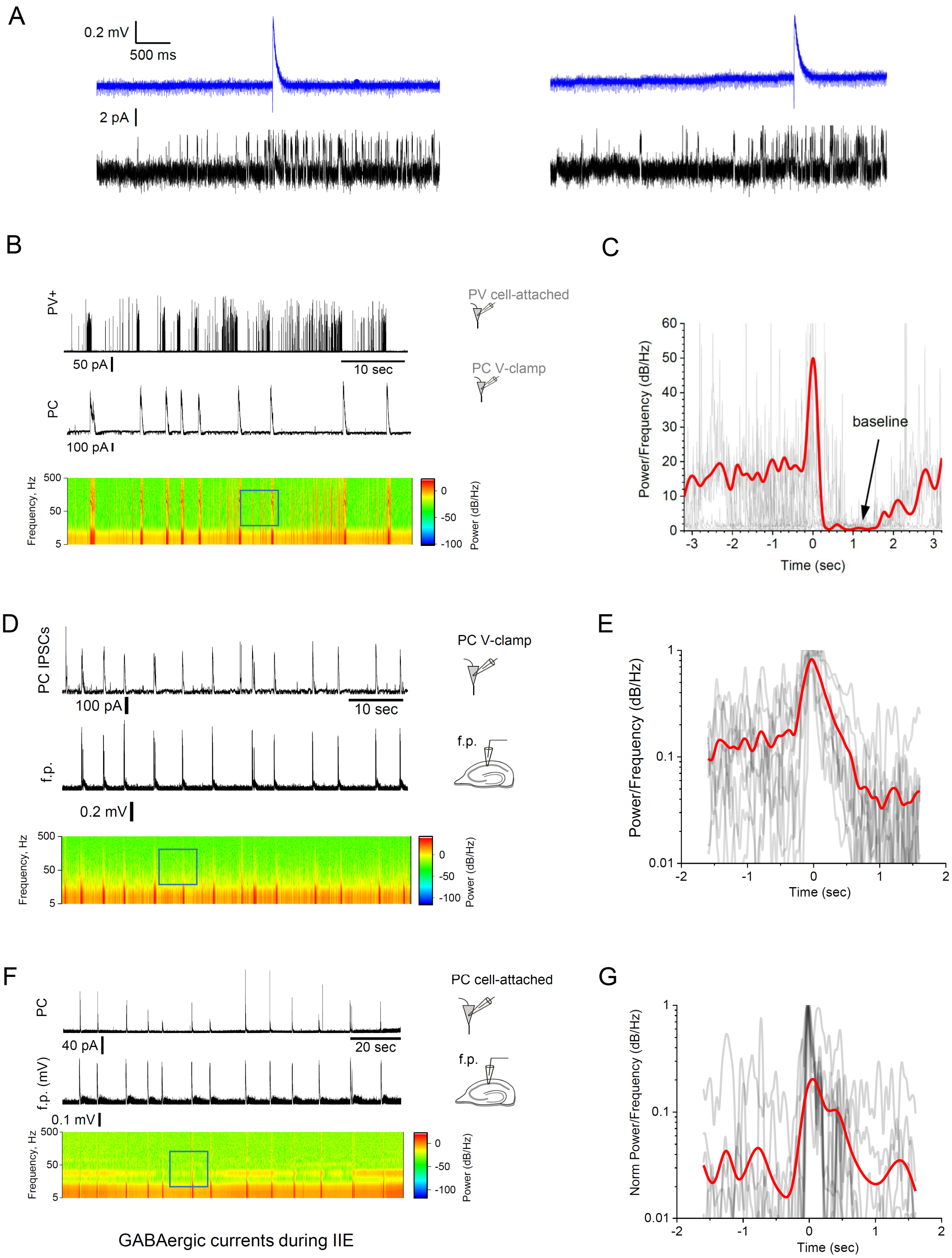


**Figure S1. PV+ interneurons increase firing rate before inter-ictal events.**

(A) Examples of local field potentials (f.p., top trace) and sniffer-patch single-channel activity (bottom) before and after individual interictal discharges in hippocampal slices.

(B) Simultaneous cell-attached spiking recording in a PV+ interneuron (top) and whole-cell recording of $GABA_A$R IPSCs in a pyramidal neuron (middle), with the power spectrum density of interneuronal spiking (bottom).

(C) Average power density across the spectrum, over the typical cycle of interictal spiking, centred at burst peak (blue square in B); red line, sample average; grey, individual interictal spike traces (n = 10, alignment at field potential spike onset, $t$ = 0); arrow, post-spike return to baseline, before an activity rise pre-spike.

(D) Simultaneous recordings of $GABA_A$R IPSCs in pyramidal cells (top) and local f.p. (middle), the power spectrum density of IPSCs recorded in pyramidal cells (bottom).

(E) Graph as as in C, but for pyramidal cell spiking shown in D (n = 9).

(F) Simultaneous cell-attached recordings of a pyramidal neuron (top), and local field potential (f.p.) during interictal events, with the power spectrum density of pyramidal cell spiking (bottom).

(G) Graph as as in C, but for pyramidal cell spiking shown in F (n = 10).

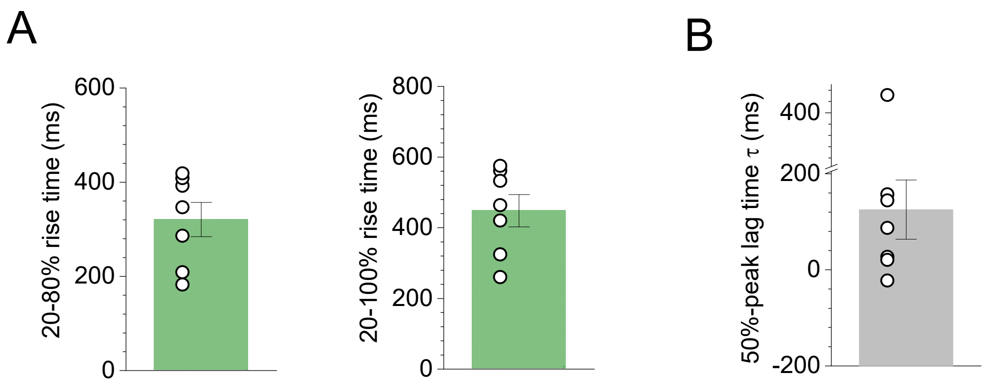


**Figure S2. Basic kinetic features of the fluorescent iGABASnFR signal that precedes interictal discharges in hippocampal slices.**

(A) Fluorescence iGABASnFR signal rise time, between 20-80% and 20-100% amplitude time points (mean ± s.e.m.: 360 ± 36 ms and 448 ± 46 ms, respectively, n = 7).

(B) Time lag between the 50%-amplitude time point of the iGABASnFR signal and the ictal fEPSP spike onset (mean ± s.e.m.: 125 ± 66 ms, n = 7).

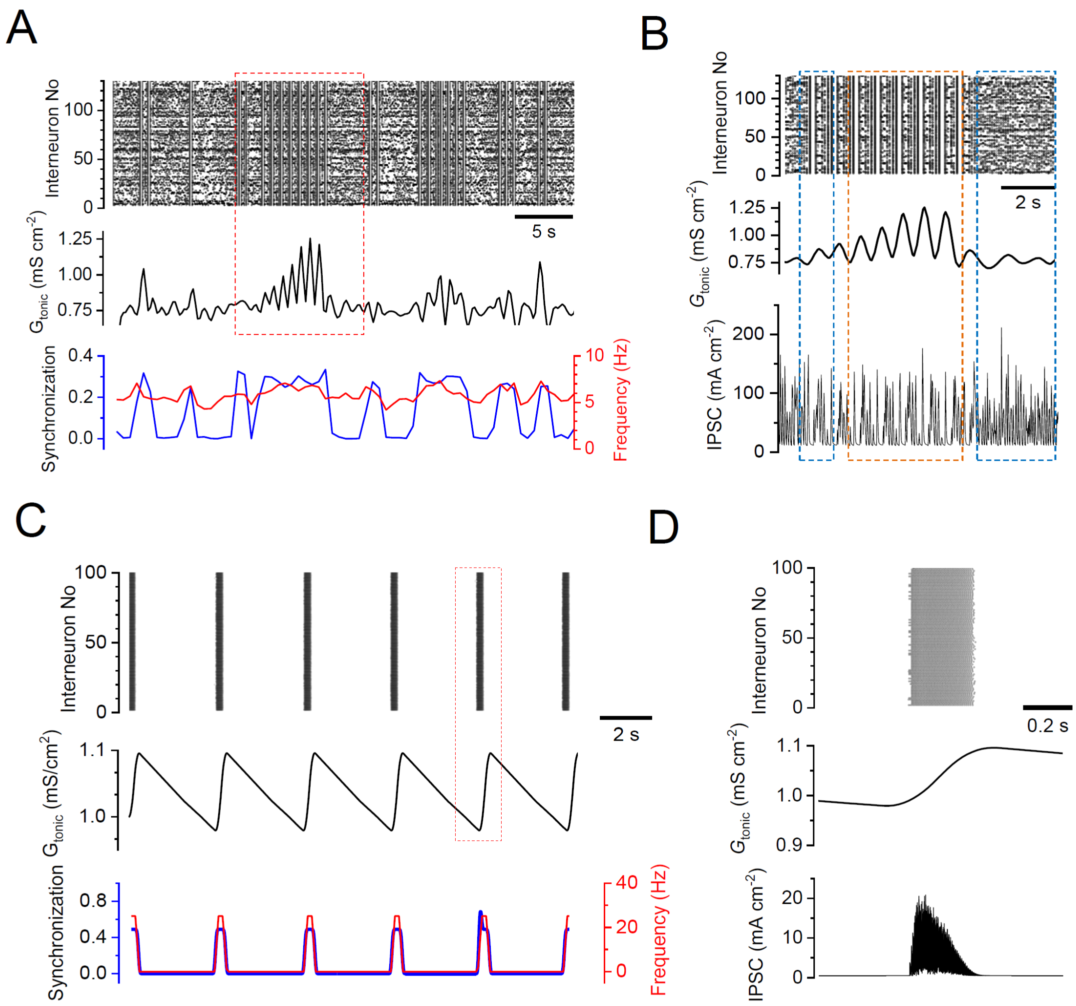


**Figure S3. [GABA]e-dependent $G_{tonic}$ drives rhythmic activity of a modelled interneuron network.**

(A) Raster plot of interneuronal network spiking activity (top), $G_{tonic}$ driven by [GABA]e (middle) calculated from integrated interneuronal discharges; and network synchrony and mean network frequency showing quasi-periodicity features (bottom). Key model parameters: cell number $N$ = 130, intra-network peak synaptic conductance $G_{ii}$ = 15 mS $cm^{-2}$; E-currents (Poisson series) with average synaptic conductance $g_s$=0.05 mS $cm^{-2}$, decay constant tau = 3 ms, and frequency $f_s$ = 50 Hz; $G_{tonic}$ reverse potential $V_{GABA}$ = –55 mV, GABA release factor $Af$ = 60 ×$10^{-7}$ nS $cm^{-2}$ $ms^{-1}$, $G_{pump}$ =0.02 $s^{-1}$ (see Methods for further detail).

(B) Fragment from A (red dotted rectangle) enlarged, with the IPSC series sampled form an arbitrarily selected simulated interneuron (bottom). Blue and orange rectangles, high-frequency non-synchronised, and oscillating and synchronised IPSC periods, respectively.

(C-D) Graphs as in A-B but with $N$ = 100, $g_s$ = 0.2 mS $cm^{-2}$ , $f_s$ = 50 Hz, $G_{ii}$ = 0.02 mS $cm^{-2}$, $V_{GABA}$ =–60 mV, $A_f$ = 0.18×$10^{-7}$ nS $cm^{-2}$ $ms^{-1}$, $G_{pump}$ = 0.0004 $s^{-1}$.

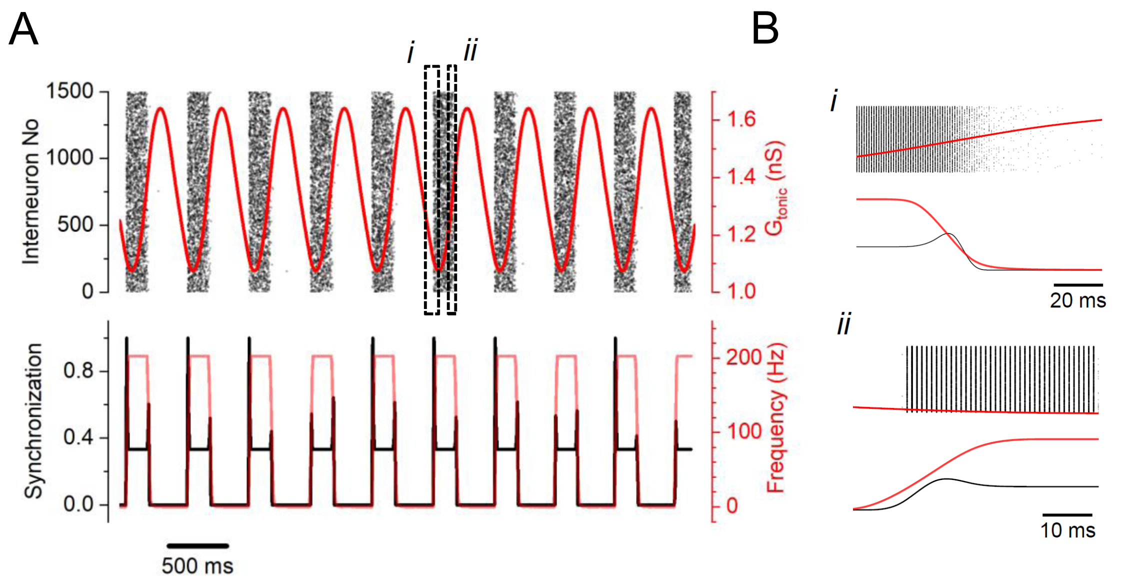


**Figure S4. Increasing network size makes burst rhythms more robust.**

(A) Raster plot of network spiking (top), network synchronisation coefficient (bottom, black), and time course of average network spiking frequency (bottom, red) for the network as in Fig. 3c; key model parameters: cell number $N$ = 1500, intra-network peak synaptic conductance $G_{ii}$ = 15 nS cm$^{-2}$; E-currents (Poisson series) with average synaptic conductance $g_s$=0.05 nS, decay constant tau = 3 ms, and frequency $f_s$ = 50 Hz; $G_{tonic}$ reverse potential $V_{GABA}$ = –55 mV, GABA release factor $Af$ = 60 $\times 10^{-7}$ nS cm$^{-2}$ ms$^{-1}$, GABA uptake rate $G_{pump}$ =0.02 s$^{-1}$ (see Methods for further detail).

(B) Fragments *i* and *ii* from A (dotted rectangles), as indicated, on an expanded scale; note that synchronization (black line, bottom traces) roughly follows the first time derivative of of frequency (red trace).

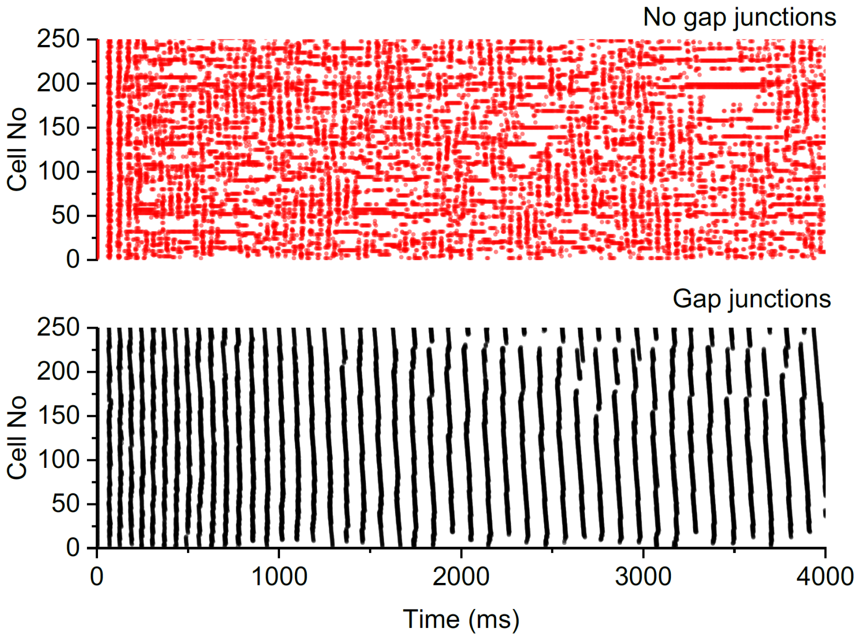


**Figure S5. Electrical coupling via gap junctions prompts high-frequency oscillations in the interneuronal network.**

Example, raster plots of interneuronal spiking (n=250 cells, model parameters as in Figure 3B), without electrical coupling (top) and with pairs of neighbouring cells connected via gap junctions (individual conductance 0.1 mS/cm$^2$; bottom).

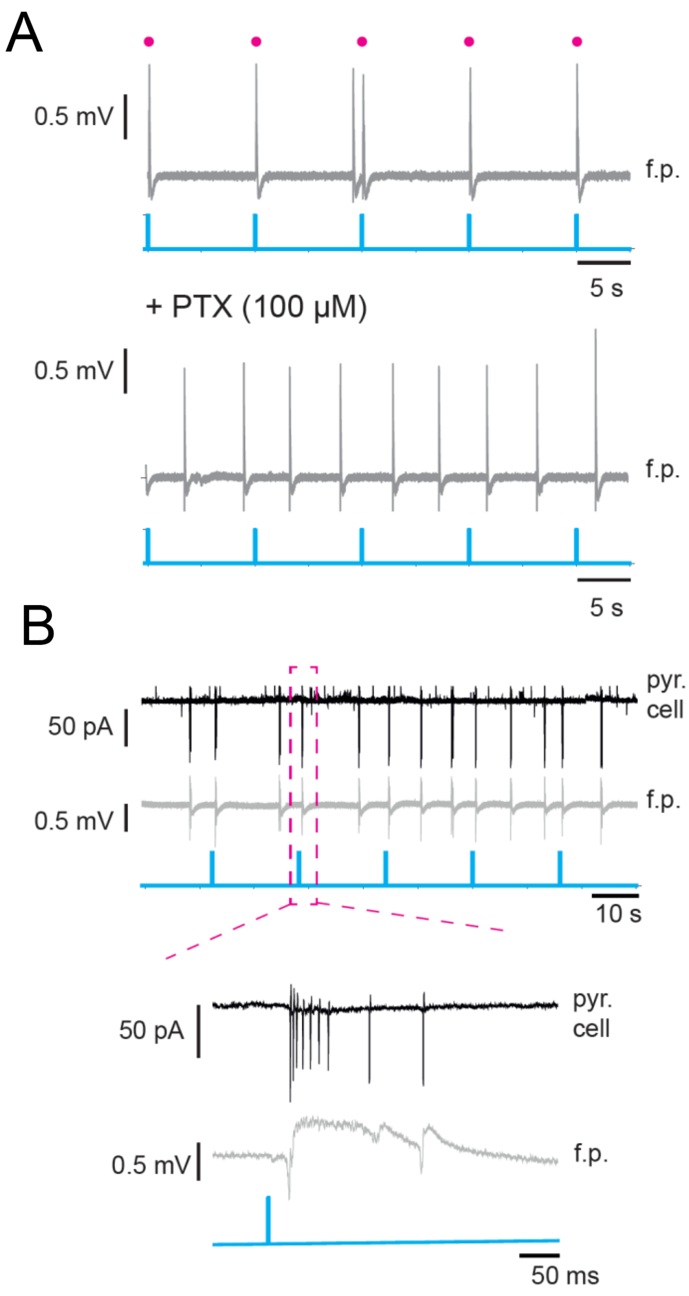


**Figure S6. Light-evoked interictal discharges are $GABA_A R$-mediated engaging principal cells**.

(A) Example of local field potential (grey trace) recorded in area CA1 (5 mM $K^+$, 0 mM $Mg^{2+}$) during optogenetic activation of PV+ interneurons (blue), with optogenetic stimulation (1ms pulses) reliably evoking interictal events with intact $GABA_A Rs$ (top panel, magenta dots), but not in picrotoxin (PTX; bottom panel).

(B) Concurrent cell-attached recording of a pyramidal neuron with the local field potential in CA1 region during interictal events.

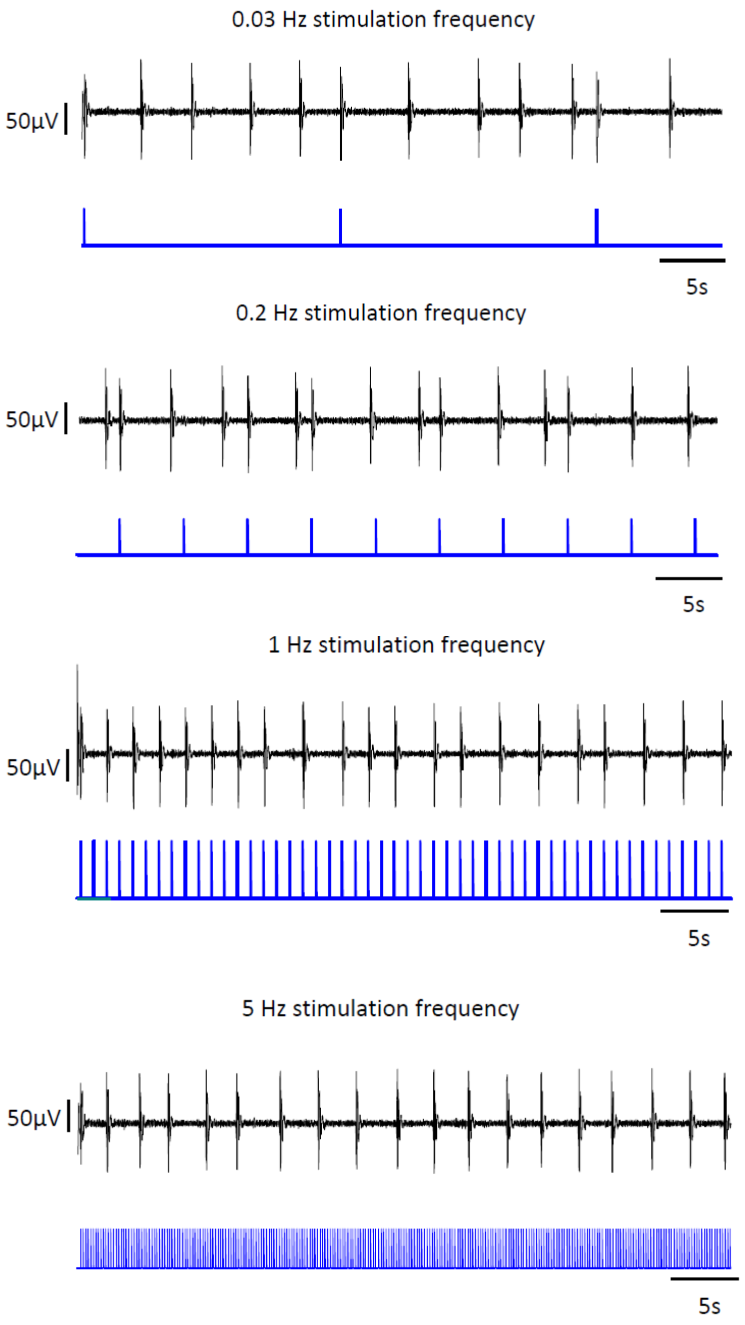


**Figure S7.** Recording of IIEs (top traces, black; bandpass filter 1-50 Hz) at varied optogenetic stimulation frequencies (bottom traces, blue; individual 1 ms pulses at 470 nm wavelength).

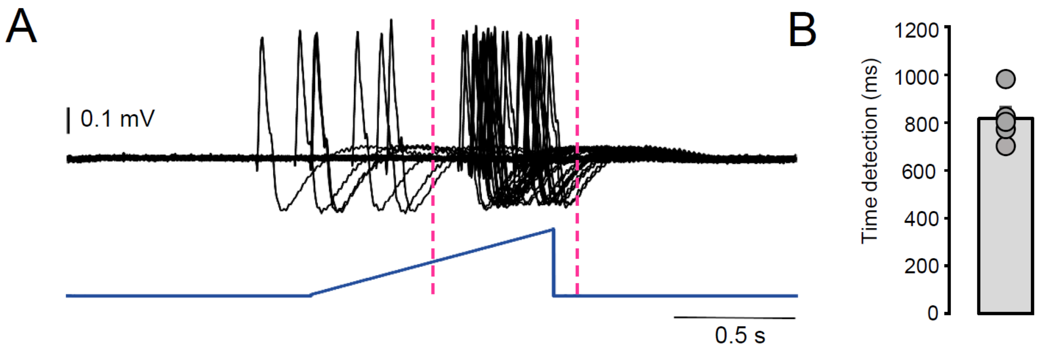


**Figure S8. Juxtaposition of the ramp stimulus and interictal discharges.**

(A) Example, superimposition of the ramped light stimulus and interictal discharges; dotted lines, 500 ms after the ramp onset, and 100 ms after the ramp endpoint.

(D) Average timepoint of the interictal spike after the ramp onset (bar, mean ± s.e.m.; dots, individual experiments; n = 5 slices in 4 animals).

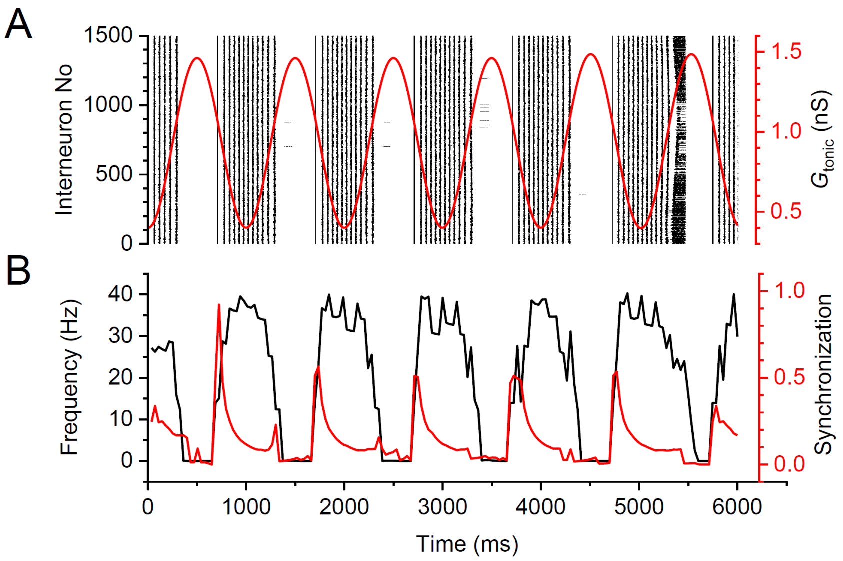


**Figure S9. Imposing cycling changes in $G_{tonic}$ drives rhythmic network activity.**

(A) Raster plot of cell spiking (black) under 1 Hz sin wave of $G_{tonic}$ (red), as indicated.

(B) Average firing frequency and synchronization parameter for network activity shown in A. Key model parameters: cell number $N$ = 1500, intra-network peak synaptic conductance $G_{ii}$ = 0.096 nS cm$^{-2}$; E-currents (Poisson series) with average synaptic conductance $g_s$ = 0.05 nS, decay constant tau = 3 ms, and frequency $f_s$ = 20 Hz; $G_{tonic}$ reverse potential $V_{GABA}$ = –53 mV, GABA release factor $A_f$ = 10$^{-8}$ nS cm$^{-2}$ ms$^{-1}$, $G_{pump}$ =0.004 s$^{-1}$ (see Methods for further detail).

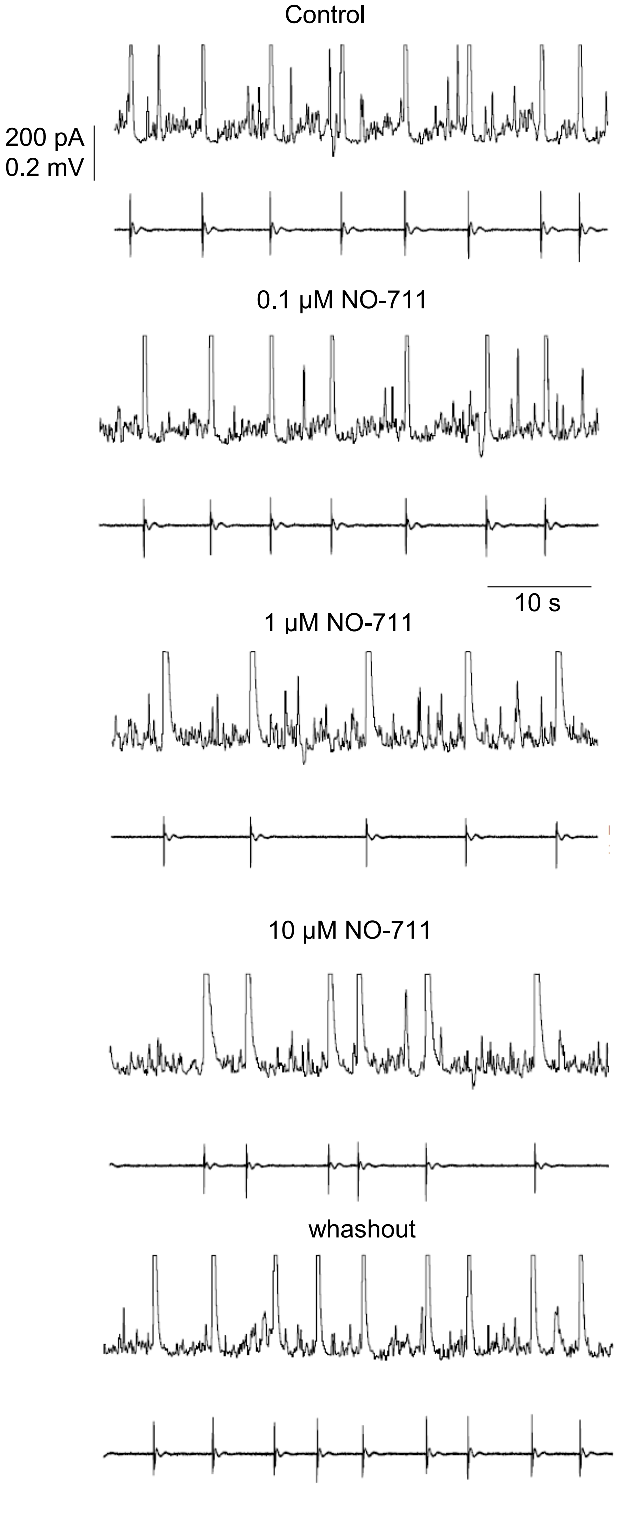


**Figure S10.** Example, simultaneous recording of CA1 pyramidal cell GABAergic currents (top traces; voltage clamp at +10 mV; 30 Hz lowpass filter) and local field potentials (bottom traces, 1-30 Hz bandpass filter), at varied concentrations of NO-711, as indicated; scales as indicated apply throughout.
